# La protein binding to telomerase RNA supports an evolutionary relationship between plant and ciliate telomerase pathways

**DOI:** 10.64898/2026.01.19.700320

**Authors:** Leon Jenner, Dzmitry Pruchkouski, Barbora Štefanovie, Olga Nováková, Monika Kubíčková, Petr Fajkus, Marie Brázdová, Jan Paleček, Eva Sýkorová

**Affiliations:** Institute of Biophysics of the Czech Academy of Sciences, Královopolská 135, 61200 Brno, Czech Republic; Laboratory of Functional Genomics and Proteomics, NCBR, Faculty of Science, Masaryk University, Kamenice 5, 62500 Brno, Czech Republic; Mendel Centre for Plant Genomics and Proteomics, Central European Institute of Technology, Masaryk University, Kamenice 5, 62500 Brno, Czech Republic; Core Facility Biomolecular Interactions and Crystallography, Central European Institute of Technology, Masaryk University, Kamenice 5, 62500 Brno, Czech Republic

## Abstract

The *Arabidopsis thaliana* La1 (AtLa1) protein is a member of the genuine La family of RNA biogenesis proteins, which are structurally similar to the La-resembling protein 7 (LARP7) family. LARP7 proteins participate in the biogenesis of the telomerase ribonucleoprotein complex in model systems, but are absent in plants. We show that AtLa1 binds to telomerase RNA in a manner reminiscent of the *Tetrahymena* LARP7 protein p65. Classical *in vitro* methods and microscale thermophoresis (MST) were used to specify the molecular structures involved in this multi-surface interaction. AtLa1 also enhances the binding of TR to the telomerase reverse transcriptase RNA binding domain. We therefore propose that biogenesis of telomerase RNA in plants and ciliates is achieved by a similar pathway, differing in the employment of genuine La or LARP7-like proteins, respectively. We also report that the domain of unknown function (DUF3223, DeCL) found in the AtLa1 protein binding partner, Domino, is an RNA binding domain with modest TR-binding capacity. This domain is also found in plant and ciliate proteins, including plant polymerases IV/V and the *Tetrahymena* La protein Mlp1. Together, these suggest that RNA biogenesis pathways in plants and ciliates have a conserved evolutionary relationship, with parallels between their La proteins.

## INTRODUCTION

The RNA-binding La protein family emerged during the early evolution of eukaryotes, characterised by a La-motif and a downstream RNA-recognition motif (RRM), which together form a La module (LaM, Figure 1A) (1–6). Substrate recognition of La proteins in most species also involves a variant RRM domain close to the C-terminus (2). Genuine La proteins typically bind RNA shortly after transcription by polymerase III, often acting as chaperones or regulators of RNA folding (1). Members of the La-related protein (LARP) superfamily are continuing to be characterised and grouped into subfamilies based on variations in structure and biological roles (2). These include nuclear LARPs that function in processing and maturation of precursor-tRNA, spliceosomal and other snRNA transcripts, and cytoplasmic LARPs involved in mRNA metabolism and translation (7,8). Recent work by Kao et al. 2024 (9) linked nuclear LARP proteins, namely LARP7 and LARP3 (a genuine La protein in humans), to sequential processing of human telomerase RNA precursors, using an *in vitro* approach. Searches of the genome of the model land plant *Arabidopsis thaliana* (At) identified eight putative La or LARP proteins (10). Of these, sequence information suggests that two are genuine La proteins, one of which (termed AtLa1, Figure 1A) can rescue non-coding RNA biogenesis functions in yeast missing their native genuine La protein, which is not the case for the AtLa2 protein (10). AtLa1 also binds pre-tRNA *via* the UUU-3′OH (poly-U) tails common to polymerase III transcripts, is located in the nucleus and is essential for embryogenesis (10). Intriguingly, the same AtLa1 protein was identified as the top co-purified protein with the full length At telomerase reverse transcriptase (TERT) in pulldown experiments (11). However, there is no direct AtLa1 protein-protein interaction with AtTERT (12). Telomerase is a ribonucleoprotein complex responsible for synthesising short repetitive DNA sequences at 3’ ends of linear chromosomes, providing ablative protection against the end replication problem in cells where it is active (13). The catalytic core of all known telomerases consists of the protein subunit TERT and a non-coding RNA, termed telomerase RNA (TR, (14,15), see (16) for review). Despite high evolutionary variability (17), TRs have conserved structural features, namely the template region and a pseudoknot (18,19); often, they also feature one or more stem-loop arms which serve as a flexible scaffold for assembly of the telomerase RNP complex (16,19).

**Figure 1:**
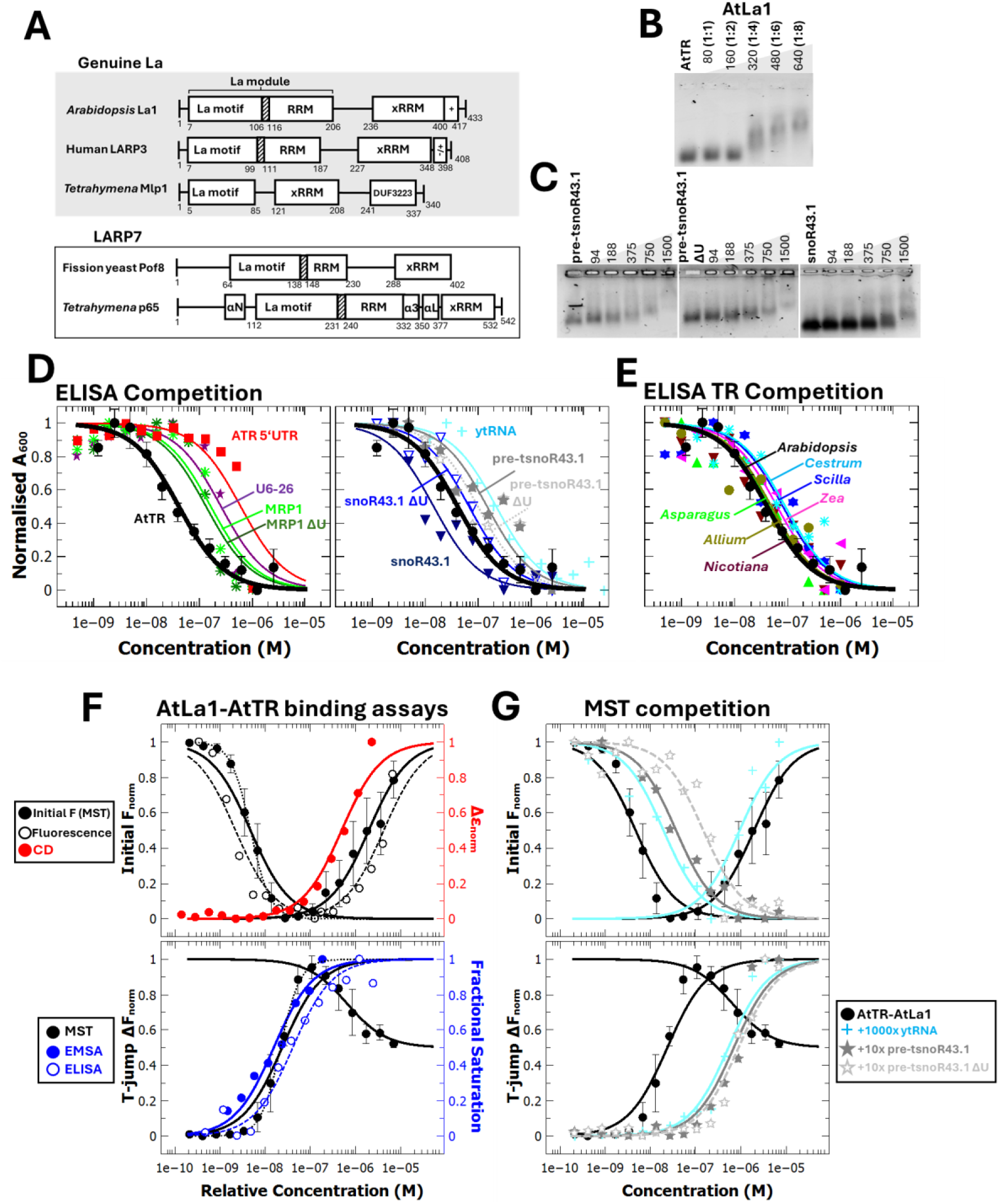
Specificity and Affinity of AtLa1 binding to AtTR. (**A**) Domain organisation of La and LARP7 proteins (2,33,37), *Tetrahymena* Mlp1 is based on Alphafold3 domain predictions and sequence alignments (Figure S5A) and includes an apparent additional xRRM in a region known to bind RNA (37). Domains are as annotated, ‘+’ or ‘-‘ indicates regions of the sequence with correspondingly charged residues at neutral pH. (**B**) AtLa1 EMSA using 1% agarose gel with 80 nM AtTR 1-268 and protein concentrations (nM) as annotated, visualized by fluorescent staining. (**C**) Representative AtLa1 EMSAs with pre-tsnoR43.1, pretsnoR43.1ΔU and snoR43.1 substrates (10,34), as annotated, conditions as (B). White lines have been added to separate panels from the same gel. (**D**) ELISA experiments detecting immunolabelled AtLa1 bound to ≤0.5 pmol biotinylated AtTR 1-268 and non-biotinylated competitor RNA as annotated, (ytRNA, budding yeast tRNA). Lines show equations fit to data used to calculate *K*_D_, average AtTR values are shown in all experiments for comparison, error bars are the standard deviation. (**E**) ELISA experiments as (D) with plant TR competitors (full details in Table S2). (**F**) Comparison of AtLa1-AtTR binding assays used in this work. Solid black circles are average initial capillary fluorescence values (upper panel) or 1.5 s T-jump data (lower panel) from MST experiments, signal is from 5 nM 3’ Cy5-labelled AtTR 1-268. Solid lines show equations fit to data used to calculate K_D_ (Table S3), dotted lines for both fast processes are Hill equation fits with n=2 for comparison, error bars are standard deviation. Open black circles with dashed lines are normalised fluorescence emission at 680 nM from 20 nM AtTR-Cy5, with excitation at 625 nM using a traditional spectrofluorometer (full spectra in Fig. S1H). Closed red circles show normalised Δε at 265 nM from circular dichroism experiments monitoring 50 nM unlabelled AtTR (difference spectra in Fig. S1I). Blue closed circles are normalised fractional saturation of 40 nM unlabelled AtTR calculated from band shift distances for EMSAs (Fig. S1J) and open blue circles and dotted lines are 1/normalised signal change for ELISA competition experiments (derived from (D) for comparison). For all experiments other than ELISAs, AtLa1 concentrations are divided by the fold increase in AtTR concentration. (**G**) MST experiments detecting AtTR-Cy5 in the presence of excess unlabelled competitor RNA as annotated (Table S4), AtTR-AtLa1 data are reproduced from (F) for comparison.

Before its role in telomerase was identified, AtTR was termed ATR8 and investigated for its role in hypoxia responses (20). AtTR is a polymerase III transcript (see (16) for review), and although there is some question of where exactly the 3’ end terminates (15,20), the short poly-U tail is tolerated for reconstitution of telomerase activity *in vitro* (14,15,21). In eukaryotes, it is typical for the *in vivo* telomerase complex to interact with multiple accessory proteins responsible for its biosynthesis and regulation (reviewed in (22,23)). Mature, catalytically-active complexes feature species-specific proteins with both parallels and differences between organisms (24,25), reviewed in (26). Proteins from the LARP7 family are one such conserved group, proposed to have a role in TR biogenesis and as subunits of the active telomerase complex, namely p65 and p43 in Alveolates *Tetrahymena* and *Euplotes* (27,28) respectively, and Pof8 (Lar7) in *Schizosaccharomyces pombe* (29–31)(Figure 1A). Telomeres progressively shorten when human LARP7 is absent (32), and this protein can counterbalance LARP3 in the processing of immature forms of human TR (9) after its transcription from the polymerase II promoter. Despite this, LARP7 is not present in active telomerase, with its role seemingly replaced by histones H2A and H2B (25). LARP7 proteins have a domain structure similar to that of genuine La proteins (Figure 1A). One difference is that their RRM2 domain, termed an extended helical RRM (xRRM), has an additional helix and conserved features that allow for a variant mode of RNA binding to specific patterns of ss- and dsRNA. Examples of this are Pof8 binding to the telomerase RNA subunit in *S. pombe* (31) and *Tetrahymena* p65 binding to its respective TR (33). Curiously, plants lack an obvious LARP7 protein, although some plants, including At, have two genuine La proteins (2,10). AtLa1 also fulfils expected genuine La functions, e.g., pre-tRNA and snoRNA binding (10,34), and is able to functionally compensate loss of budding yeast homolog Lhp1p *in vivo*, unlike the AtLa2 protein (10). It is therefore plausible that the co-purification of AtLa1 observed in a TAP/MS study (11) may be caused by its interaction with AtTR and/or with other co-purified telomerase partners.

AtLa1 has a single known protein-protein interaction with Domino (12,35), a plant-specific protein of unknown function which is located in the nucleus, is essential for embryogenesis (36), co-purifies and interacts with AtTERT (11,12). Intriguingly, the domain of unknown function (DUF3223, aka. DeCL) in Domino is also found in the genuine La proteins of Alveolates (2,37), (Figure 1A) as well as being present in the C-terminus of plant-specific AtPolymerase IV and V (38,39), and defective chloroplasts and leaves (DCL) proteins (40). The At telomerase complex to date lacks full characterisation, although binding between specific domains of AtTERT and AtTR has been characterised (21), and an interaction between AtTR and AtNAP57 (aka. Cbf5 or dyskerin) was identified (41). Plant and ciliate TR genes share structural and regulatory characteristics that suggest they are transcribed by RNA polymerase III (14). The recent discovery of plant-type TRs in insects (42) enlarge the number of evolutionarily distant groups sharing the same pathways of TR biogenesis (see (16) and references herein). It is therefore of interest to investigate the specificity of AtLa1 binding to AtTR, including whether such binding is part of telomerase-specific maturation or even if AtLa1 might be a component of the active telomerase complex, given the structural similarity between La and LARP7 protein families. This study determines structural regions of both molecules responsible for AtLa1-TR binding and their affinity. Of secondary interest is the structural basis of AtLa1 binding AtDomino, and if there is an obvious function for the DUF3223 motif, so far found only in plants and ciliates (2,36–40).

## MATERIAL AND METHODS

### Preparation of protein and RNA constructs

Preparation of Gateway entry clones and Gateway Y2H destination clones coding for the full-length AtLA1 (AT4G32720) and DOMINO1 (AT5G62440) was described previously (12). Constructs for the AtTERT (AT5G16850) TRBD domain for Y3H were prepared previously (21). Constructs bearing variant AtLa1 and AtDomino fragments were subcloned using specific primers (Table S1). Constructs bearing mutated regions of AtLa1 (F331A, D333R, R344A, W387A, SYD124GGG) and AtTR (gg179cc, ggg179cac, gg185cc, gg193cc, gg196cc) were prepared by mutagenesis of AtLa1 plasmids and AtTR plasmid, respectively using a QuikChange Lightning Site-Directed Mutagenesis Kit (Agilent Technologies, #210519) according to the manufacturer’s instructions (all primers are listed in Table S1).

Plasmids bearing plant TR constructs of *A. thaliana*, *Scilla peruviana*, *N. sylvestris*, *Allium cepa* and control construct pCDT1a were prepared previously (14). RNA constructs coding for snRNA MRP1, U6.26, pre-tRNA-snoR43.1 and the 5’UTR ATR sequence were amplified from genomic DNA of *A. thaliana* (Col-0) with specific primers (Table S1) and PCR products were introduced to pCR II TOPO vector (Invitrogen, #K461020). Specific primers for cloning of plant TR constructs were designed using assembled TR subunit sequences of *Pisum sativum*, *Solanum lycopersicum*, *Cestrum elegans*, *Zea mays*, *Asparagus officinalis* published in (14). Plant TR sequences were amplified using genomic DNA isolated from leaves of *C. elegans* L. (43) or young seedlings of *Z. mays* L., *P. sativum* L., *S. lycopersicum* L. and *A. officinalis* L., seeds were purchased from local distributors (Nohel Garden, SEMO Smržice). PCR products were introduced to pCRII TOPO vector (Invitrogen, #K461020) and plasmids were sequenced to determine the construct orientation in respect to the universal M13F primer. The AtTR constructs ΔP4, mP2/3 were prepared previously (21), ΔT/PK (deletion nt 19-171) and P4/5/6 (nt 178-251) constructs were prepared using specific primers listed in Table S1. The ΔP6 (deletion nt 201-231) construct was generated using PCR product amplified with specific primers (Table S1), AtTR plasmid as a template and Phusion^®^ High-Fidelity DNA Polymerase (Thermo Fisher Scientific, #F530L). The ΔP6 plasmid was recovered using PCR product and GeneArt™ Gibson Assembly HiFi Master Mix (Thermo Fisher Scientific, #A46624) according to the manufacturers’ instructions, mutation was verified by plasmid sequencing. AtTR fragments ΔT/PK, ΔP6 and P4/5/6 for Y3H were PCR amplified (primers specified in Table S1) and ligated into the SmaI site of pIIIA/MS2-2 vector using NEBuilder® HiFi DNA Assembly Master Mix (New England Biolabs, #E2621S).

### Protein expression and purification

The Gateway cloning system (Invitrogen) was used to introduce DNA fragments encoding desired proteins and protein fragments into pDEST15 or pDEST17 Gateway destination vectors according to the manufacturer’s instructions. These constructs allow for the expression of proteins with N-terminal glutathione transferase (GST) or poly-histidine (His) tags for affinity purification. Plasmids containing protein constructs were then transformed into Rosetta (DE3) pLysS or BL21-CodonPlus-RIL cells. 0.5 mM to 0.6 mM isopropyl-1-thiol-b-D-galactopyranoside was added to the log-phase cell culture to induce protein expression overnight at 20 to 25 °C. Cells were harvested at 5000 rpm at 4 °C for 15 min and either stored at -80 °C for later use or immediately disrupted by sonication in lysis buffer (0.1 M Tris-HCl pH 8.0, 0.5 M NaCl, 0.5% w/v polyethyleneimine (SERVA, #3314103), 1mMbenzamidine-HCl (Carl Roth, #R.CN38.3), 1 mM polymethyl sulfonyl fluoride, with purifications for EMSAs also including 0.5 mM dithiothreitol). Afterwards disrupted cells were centrifuged at 20,000 rpm for 10 minutes at 4 °C. The supernatant was applied to SpinTrap GST or Ni Sepharose spin columns (Cytiva, #28-4013-53, #28-9523-59), reused and periodically refilled with 0.4 mL 50% Glutathione Sepharose® 4 Fast Flow (Cytiva,# 17-5132-01) or Ni-NTA agarose Fast Flow resin (Cube Biotech, # 31105). Purification proceeded according to manufacturer’s instructions, modified to include three equilibration steps and three washing steps overall. The target protein was then eluted with 0.1 M Tris-HCl pH 8.0, 0.5 M NaCl, with the addition of 40 to 80 mM glutathione or 100 mM imidazole, for GST and His purifications respectively, and 10% glycerol, 1mM ethylenediaminetetraacetic acid and 0.5 mM dithiothreitol added to this and subsequent buffers for EMSA experiments. Imidazole or glutathione was then reduced to <1 mM concentration by buffer exchange using Amicon® Ultra Centrifugal Filters (Millipore, Merck) and the protein was concentrated as appropriate. La1 and TRBD rapidly lose binding ability over time or with freeze-thaw cycles, so all experiments were performed using proteins purified earlier the same day. Domino stocks were stored at -80 °C between experiments. Protein concentrations are estimated values based on 280 nm absorbance measured using a NanoDrop ND-1000 spectrophotometer and extinction coefficient values from *in silico* prediction (ExPaSy ProtParam tool).

### RNA synthesis and labelling

RNA was synthesised using HiScribe™ T7 Quick High Yield RNA Synthesis Kit (New England Biolabs, #E2050S) to transcribe a PCR product amplified from pCRII TOPO plasmid DNA sequences using primers detailed in (14,21) and in Table S1. AtTR, AtTR fragments and other RNA templates (MRP1, U6-26, ATR5’UTR, pre-tRNA-snoR43.1, snoR43.1, pCDT1a) were produced using PCR products prepared using T7-specific forward and specific reverse primers. For competitive ELISAs, plant TRs were transcribed using a PCR product generated from plasmid DNA using universal M13F and sequence specific reverse primers (Table S1). RNA was then purified using AMPure XP magnetic beads (Beckman Coulter, #A63881) according to the manufacturer’s protocol, eluted in water, and stored at -80 °C between experiments. For ELISA or MST experiments, synthesised RNA was 3’ labelled with biotin- or Cy5-labelled cytosine bisphosphate, respectively (Thermo Scientific #20160, Jena Biosciences #NU-1706-CY5), using Pierce™ RNA 3’ End Biotinylation Kit (Thermo Scientific, #20160) according to the manufacturer’s instructions. Labelled RNA was stored away from light at - 80 °C. Alternatively, 5’ Cy5-labelled RNA was instead prepared using a 5’ End Tag™ kit (Vector Laboratories, #MB-9001) to de-phosphorylate RNA and re-phosphorylate with a γ-[S] phosphate group, followed by covalent linkage of S to Cy5-maleimide (Vector Laboratories, # FP-1322-1). The manufacturer’s instructions were followed with the exception that the sulfur-maleimide reaction was performed overnight. Efficiency of labelling by this method was generally poor (ca. 10%) following the instructions directly, this was optimised to 60% following overnight maleimide linking, but remained consistently lower efficiency than 3’ labelling (≥80% efficiency consistently) in our hands (details in Supplementary information). For northwestern blotting (Supplementary data and Figure S1), synthesised RNA was dephosphorylated using incubation with quick calf intestinal phosphatase (Quick CIP #M0525, New England BioLabs) for 1 hour at 37 °C, before purification using AMPure XP magnetic beads (Beckman Coulter, #A63881) or High Pure RNA isolation kit (Roche, #11828665001) according to the manufacturer’s instructions. Dephosphorylated RNA was then re-phosphorylated with radioactive ^32^P using incubation with γ-[^32^P] ATP and T4 polynucleotide kinase (Thermo Scientific, #EK0032) incubation for 1 hour at 37 °C followed by purification using MoBiSpin G-50 gel columns (MoBiTech GmbH, #SCO510). Radiolabelled RNA was then stored at -20 °C until use. Prior to all experiments, RNA stocks were heated at 95 °C for 5 minutes before immediately being placed on ice to refold.

### Electrophoretic mobility shift assays

*In vitro* synthesised RNA was mixed with protein fragments and variants of specified concentrations in 20 uL of reaction buffer per sample, with final concentrations of 0.1 M Tris-HCl pH 8.0, 0.5 M NaCl, 10% Glycerol, 1mM Benzamidine-HCl, 1mM EDTA, 1 mM PMSF, 0.5mM DTT and typically 80 nM RNA concentration. The mixtures were then incubated on ice for 20 min before the protein-RNA complex was resolved by 1% non-denaturing agarose gel in 1x TAE buffer at 4 °C, 50 minutes, 90 V. Gels were stained with 5000x diluted EliDNA PS Green (Elisabeth Pharmacon, Czech Republic) in 1x standard TAE buffer at room temperature for 30 minutes and then imaged using an Amersham™ Imager 680 (GE Healthcare). Gel images are representative examples of experiments repeated at least 3 times. For La1 WT, the mean displacement of bands from two EMSA repeats using a larger concentration range was measured from the relative position of overlaid lines in Microsoft PowerPoint and further analysed in Microsoft Excel and QTI plot (see below for *K*_D_ determination).

### Enzyme-linked immunosorbent assays

96-well Immuno plates (SPL Life Sciences) were incubated overnight with either 5 μg/mL streptavidin (Sigma Aldrich, #S4762) at room temperature or 5 μM purified AtLa1 His at 4 °C, 50 μL per well in 1x PBS. Plates were washed with 1x phosphate-buffered saline (PBS) and then blocked with 3% bovine serum albumin (Sigma Aldrich, #A3294) for 2-3 hours at room temperature. For experiments with RNA, plates were then washed with 1x PBS and then incubated for 1-2 hours with 4 nM 3’ biotinylated AtTR 1-268 in 5 mM Tris-HCl, 50 mM KCl, 0.01% Triton X-100, pH 7.6. Plates were washed with 1x PBS with 0.05% Tween (PBST), then incubated with purified GST fusion proteins, pre-incubated with 1:2 anti-GST antibodies (G1160, Sigma Aldrich), in a range of concentrations as appropriate in 100 mM Tris-HCl, 500 mM NaCl, pH 8.0 for 20 minutes. Plates were then washed with 1x PBST and then incubated with 0.5% BSA in 1x PBS for 5 minutes three times, before being incubated with 1:1000 secondary anti-mouse antibodies (A0168, Sigma Aldrich) washed and blocked a further three times in the same way, and then developed using 0.1 mg/mL tetramethylbenzidine (Sigma Aldrich, #860336) and 0.04% H_2_O_2_ in 50 mM phospho-citrate buffer pH 5.0 for 5 minutes. Absorbance at 600 nm was then recorded for each well twice using a synergy H1 plate reader (Agilent BioTEK). At least two independent repeats of each experiment were performed, with absorbance values normalised, curated and averaged using Microsoft Excel and QTIplot (IonDev Software, see below for *K*_D_ determination).

### Microscale thermophoresis

2-fold serial dilutions of GST fusion proteins were mixed in a 1:1 ratio with Cy5-labelled RNA to give final concentrations of typically 0.2-10,000 nM protein and 5 nM RNA in 50 mM Tris-HCl, 250 mM NaCl, 250 mM potassium acetate, pH 7.0 with 0.05% Tween. Sample preparation took approximately 5 minutes before samples were loaded into normal or premium capillaries (NanoTemper) and both initial fluorescence and microscale thermophoresis (MST) measurements for up to 20 seconds were recorded using 1-8% power laser excitation and a Monolith NT.115 pico instrument (NanoTemper),. Raw data were exported and analysed using Microsoft Excel and QTIplot (IonDev Software, see below for *K*_D_ determination). All MST values presented are those recorded at 25 °C from the 1.5 second T-jump as in many cases aggregation caused high levels of noise in longer MST traces.

### *K*_D_ determination

Change of absorbance (ELISA), band displacement (EMSA), circular dichroism (CD) or fluorescence (MST) data were normalised and then fit to the rearrangement of the standard equation for dissociation constant (*K*_D_) determination:

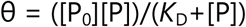

Where θ is fractional saturation (*ie.* the normalised absorbance or fluorescence signal), [P] is protein concentration and [P_0_] is protein concentration in the absence of RNA (defined as 1 or -1 after normalisation, depending on the direction of the process). Fittings were performed using QTIplot (IonDev Software) and a scaled Levenberg-Marquardt algorithm for non-linear least-squares fitting. Numbers of replicas for ELISA and MST experiments are given in Tables S2, S3 and S4.

### Yeast two-hybrid assay

The classical Gal4-based Y2H system was used to analyse protein-protein interactions as described previously (44). Briefly, pGBKT7 and pGADT7 constructs were co-transformed into the S. cerevisiae PJ69–4a strain and selected on SD -Leu, -Trp plates. Drop tests were carried out on SD -Leu, -Trp, -His (with 0.1; 0.5; 1; 5; 10; 15; 30 mM 3-aminotriazole) plates at 28 °C. Each combination was co-transformed at least three times, and three independent drop tests were carried out.

### β-galactosidase assay

Assay was performed as described in (21). After the transformation of the ∼500 ng of each plasmid DNA, single colonies were inoculated into 100 µL of YPD medium (in triplicates) in a 96-well plate and grown overnight at 28 °C. The next day, OD_600_ was measured using a BioTEK Powerwave 340 microplate reader (Agilent BioTEK) by diluting 10 µL culture in 90 µL water. The remaining 90 µL of the cultures were washed in 50 µL of Z-buffer (60 mM Na_2_HPO_4_, 60 mM NaH_2_PO_4_, 10 mM KCl, 1 mM MgSO_4_, pH 7.0), 25 µL of 0.1% SDS was added, followed by addition of 6 µL of chloroform. Reactions were mixed several times by pipetting and incubated for 15 min at 30 °C. Next, 60 µL of 4 mg/mL ortho-Nitrophenyl-β-galactoside (Sigma Aldrich, #N1127) was added to each reaction and mixed well. The time of the reaction was recorded. Once the yellow colour appeared, the reactions were stopped by adding 120 µL of 1 M Na_2_CO_3_ and mixed well. The plate was spun at 2500 rpm for 2 min to remove cell debris. 100 µL of the supernatants were transferred to a new plate and OD_420_ and OD_550_ were measured using a microplate reader. Miller units were calculated according to the formula:

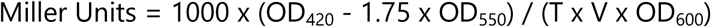

OD_420_ and OD_550_ are read from the reaction mixture, OD_600_ reflects cell density in the cell suspension, T = time of the reaction in minutes, and V = volume of culture used in the assay in mL. Relative growth in Miller units was then normalised to the highest value (AtLa1 WT), analysed and presented using Microsoft Excel and QTIplot (IonDev Software).

### Yeast three-hybrid assay (Y3H)

The Y3H system (45) was used to map protein-TR interactions as described previously (21). Briefly, variant AtTR-MS2 (pIIIA/MS2-2 derived) plasmids and Gal4AD-AtTERT fragments (in pGADT7 derived plasmids) were transformed into *Saccharomyces cerevisiae* strain YBZ-1 and selected plates missing leucine, and adenine (-LA). RNA-protein interaction was monitored by YBZ-1 cell growth on selection plates missing leucine, adenine, and histidine (-LAH). In addition, increasing concentration of 3-amino triazole (3AT), which competitively inhibits the imidazole glycerol-phosphate dehydratase (product of HIS3 gene), was used to measure the relative strength of the interaction. Each combination was co-transformed two times, and two independent drop tests were carried out.

A multicomponent variant of the Y3H assay (derived from the multi-component Y2H system; (46))was used to test the effect of La1 on the AtTERT-AtTR interaction. AtTR-MS2, Gal4AD-AtTERT and Gal4BD-La1 (pGBKT7 derived) plasmids were co-transformed into YBZ-1 strain and selected on -LWA plates. Empty Gal4AD and Gal4BD plasmids were used as a control. RNA-protein interaction was monitored on -LWAH plates with or without an increasing concentration of 3-aminotriazole (3AT). Each combination was co-transformed two times, and two independent drop tests were carried out.

### Mass photometry

Samples for mass photometry using a TwoMP instrument (Refeyn, Oxford UK) were prepared in higher concentrations (AtLa1 100 nM, 1000 nM Domino) in commercial culture grade Dulbecco’s phosphate-buffered saline (Gibco). Samples and buffers were equilibrated to room temperature before measurement, then diluted 20x typically in the final drop. Samples with AtTR (except Domino+AtTR) were instead prepared at 10 μM of all components and a pipette tip was used to add a tiny amount of sample to an 18 μL drop to catch larger complexes before they dissociate. Drops were cast on MassGlass UC (Refeyn) precast microscope slides after sonication in 50% isopropanol followed by washing with milli-Q water to maximise glass cleanliness. Mass values were calibrated against MassFerence P1 (Refeyn) and an in-house 40 kDa protein standard, with mass peaks typically giving a linear fit with R^2^ = 1.000, gradient -36.62 mDa, maximum mass error of 2.1% and intercept (contrast) of -0.00007. 60 s videos were recorded using AquireMP 2024 R2 software (Refeyn). Events recorded during this time were deconvoluted into mass histograms with 5 kDa-width bars and manually fit to Gaussian profiles using DiscoverMP v2024 R2 software (Refeyn). Following optimisation, all measurements were performed in at least triplicate, with representative results shown in Figure 5E.

## RESULTS

### La1 binds to snRNA, with preference for telomerase RNA

RNA binding experiments were performed with several small nuclear RNAs (snRNA) and other RNA constructs with structural characteristics that could be relevant to protein-RNA interactions. Work started with AtTR, given that AtLa1 copurifies with telomerase (11) but does not bind AtTERT directly (12), which would be rationalised if AtTR binds to AtLa1 as well as AtTERT. Incubating purified full-length (FL) AtLa1 protein (Figure S1A, S1B) with *in vitro* synthesised AtTR causes a shift in RNA mobility detectable by EMSA (Figure 1B) consistent with complex formation. Northwestern hybridization of immobilized proteins with AtTR confirms that the species responsible for this has the predicted mass of AtLa1 and this signal was also detected in an excess of unlabelled competitor RNA (Figure S1C). Binding of AtTR to AtLa1 occurs regardless of the presence of an N-terminal 6xHis sequence or glutathione transferase (GST) fusion used for affinity purification (Figure S1C). Consistent with this, GST alone does not cause any observable shift in EMSA experiments (Figure S2). Prior reports suggest a more general role for AtLa1 in biogenesis and processing of polymerase III transcripts (10,34). Comparable shifts in EMSAs are also observed when AtLa1 is incubated with *in vitro* synthesized variants of the dicistronic pre-tsnoR43.1 and snoR43.1 transcript (Figure 1C), or the snRNA subunit of the RNase for mitochondrial RNA processing (MRP1, Figure S3A, (47)). Curiously, AtLa1 also binds a pre-tsnoR43.1 construct missing its 3’ poly-U trailer (pre-tsnoR43.1 ΔU, Figure 1C) with similar affinity, complicating the initial assumption that AtLa1 is sensitive only to 3’ UUU-OH motifs for binding (10). Aside from polymerase III transcripts, AtLa1 binding was also detected with RNA derived from the 5’-UTR of the ATAXIA TELANGIECTASIA-MUTATED AND RAD3-RELATED (ATR) mRNA ((48), Figure S3A). This construct was tested as RNA-binding proteins often prefer G-rich regions that attract nonspecific binding, and ATR 5’-UTR is known to form stable RNA guanine quadruplexes (G4s), which have been regularly implicated in structure-specific protein-RNA interactions (49,50). G4s are known to occur in human TR ((51–53), reviewed in (54)) suggesting a precedent for G4-protein interactions in telomerase. However, it is worth noting that in the buffer used for the majority of this study, AtTR does not evidence a hyperchromic fluorescent signal in the presence of G4-reactive N-methylmesoporphyrin (NMM, (55), unlike ATR 5’UTR (Figure S1E). Thus, there is no evidence that AtTR forms G4 structures during the experiments reported here.

To investigate the preferences and specificity of AtLa1-RNA binding, competitive ELISA experiments were performed using immobilised 3’ biotinylated AtTR incubated with immunolabelled La1-GST and various mobile competitor RNAs, including those tested in previous EMSA experiments (Figure 1D, Table S2). Of these, unlabelled AtTR competes with AtLa1-AtTR binding with a higher affinity (*K*_D_ = 41 nM) than all but snoR43.1 (*K*_D_ = 14 nM). The 5’UTR ATR competes with the lowest affinity (Figure 1C, *K*_D_ = 600 nM), followed by polymerase III transcripts U6-26 (*K*_D_ = 260 nM), mature budding yeast tRNA (K_D_ = 220 nM) and RNase MRP subunit MRP1 (*K*_D_ = 160 nM). Interestingly MRP1 binds with essentially identical affinity when the polyU tail is not present (MRP1 ΔU, *K*_D_ = 130 nM), and the same is true for pre-tsnoR43.1 and pre-tsnoR43.1 ΔU (K_D_ = 150 nM and K_D_ = 100 nM, respectively). It is of interest to note the difference between the affinity of snoR43.1 and snoR43.1 ΔU, where the latter loses nearly 5-fold affinity for AtLa1 (K_D_ = 70 nM) in competitive ELISAs. This difference in affinity might be the molecular basis of the two pathways where AtLa is proposed to be part of processing the dicistronic pre-tsnoR43.1 transcript to mature snoR43.1 (34).

AtLa1 clearly has affinity for multiple pol-III transcripts *in vitro*, consistent with a flexible biological role in pol-III transcript biogenesis. However, its high affinity for TR raises the question of whether it has an additional role in the maturation of telomerase in *A. thaliana*, or indeed for plants in general. Plant TRs share similar structural characteristics, therefore to examine the possibility of a more general mode of AtLa-TR interaction, competitive ELISA RNA-protein binding experiments were performed with AtTR and nine plant TR constructs (Figure 1E). These constructs included representatives of species with variant telomeric repeats (17). AtLa1 binds all plant TRs tested with similar affinities regardless of telomere type and evolutionary relationship (Table S2). Binding at most has a 2- to 3-fold variability for TRs of *Arabidopsis thaliana* (Brassicales*, K*_D_ = 41 nM), garden pea (*Pisum sativum*, Fabales, *K*_D_ = 80 nM) and *Cestrum elegans* (Solanales, *K*_D_ = 90 nM), the latter which synthesises variant telomeric repeats. The same is true for other representatives of species with telomere variants, such as monocots maize (*Zea mays,* Poales, *K*_D_ = 60 nM), *Scilla peruviana* (Asparagales, *K*_D_ = 80 nM), asparagus (*Asparagus officinalis,* Asparagales, *K*_D_ = 50 nM) and onion (*Allium cepa,* Asparagales, *K*_D_ = 50 nM). *Nicotiana sylvestris* (Solanales, *K*_D_ = 42 nM) also has a comparable affinity, and the same for for tomato (*Solanum lycopersicum*, *K*_D_ = 30 nM). This is therefore consistent with a hypothesis that there is a general conserved role of La proteins in plant telomerases. This interaction might be simply explained if there is a sequence or structure in TRs which forms a conserved interface with one or more RNA binding domains of plant La proteins. Conserved or covariant regions in Angiosperm TRs include the pseudoknot-forming P2 and P3 regions, closing P1a stem and parts of the P4/5/6 stem (15).

### AtLa1 binding to AtTR is a dynamic biphasic process

To investigate the molecular basis of this La-TR interaction, which presumably might be generally applicable to land plants, *in vitro* thermodynamic protein-RNA binding experiments were performed. AtTR binding to AtLa1 was investigated in greater detail using microscale thermophoresis (MST, Figure 1F, Table S3) and we cross-validated the performance of MST for investigation of RNA-protein binding. The applicability of this relatively new technique (reviewed in (56)) for the detection of specific protein binding to structured RNA was first explored using AtLa1 and a known pre-tRNA substrate (10)(Figure S4A), followed by AtTR (Figure 1F, S1K). MST was also employed in competition assays (Figure 1G). As another proof of concept, MST detection of AtTR binding to its natural target, the telomerase RNA binding domain (TRBD, (21)(see Supplementary information, Figure S4C), and other comparative analyses using classical and modern techniques were performed (see Figures 1F, S1F-J). We also compared the performance of 5’- and 3’-attached fluorescent label (see Supplementary information and Figure S4D, S4E), as a consequence of which we report all other results with 3’ Cy5-labelled RNAs. In summary, the MST experiments performed here reveal data on protein-RNA interactions in two ways (demonstrated in Figure 1F). Firstly, via initial fluorescence changes of an AtTR 3’-attached Cy5 fluorescent label, which is sensitive to its environment (upper panel, Figure 1F, Figure S1F-G), reporting on structural changes from interactions as a biphasic process comparable to classical fluorescence titrations (open circles, Figure 1F, Figure S1H). The fast process (K_D_ = 5 ± 1 nM) is one of protein-induced fluorescence quenching (PIFQ, (57)) likely to report on smaller-scale structural changes within the region of the Cy5 label. The slow process (K_D_ = 1.9 ± 0.2 μM) is a protein-induced fluorescence enhancement (PIFE, (58)) that occurs concomitantly with perturbation of the RNA A-type helical structure monitored by circular dichroism (red circles, Figure 1F, Figure S1J).

Thermophoresis proper is also recorded, in this case initial changes in the first 1.5 seconds of the experiment, known as the temperature jump (T-jump, Figure S1G). This process reports on apparent mass changes resulting from *bona fide* protein-RNA complex formation (lower panel, Figure. 1F), which is also biphasic. The fast T-jump process (K_D_ = 20 ± 10 nM) is consistent with apparent mass changes in EMSAs (blue circles, Figure 1F), displacement of the complex with competitor RNA in ELISAs (open blue circles, Figure 1F) and even sedimentation of the AtTR-La1 complex (Figure S4G). The second process is often incomplete or unresolved over the concentration ranges used for binding experiments (Table 1). We will not consider the latter process further, but initial fluorescence fast and slow processes will be compared and considered as relating to mostly specific interactions, and the fast T-jump process will be considered as reporting on overall binding, including non-specific contributions similar to ELISA or EMSA techniques.

**Table 1.** *K*_D_ values calculated from MST experiments at 25 °C.

| La1 | AtTR | Fluor 1 [nM] | Fluor 2 [μM] | MST 1 [nM] | MST 2 [μM] |
| --- | --- | --- | --- | --- | --- |
| **FL** | **1-268** | $5 \pm 1$ | $1.9 \pm 0.2$ | $20 \pm 10$ | $0.65 \pm 0.1$ |
| | **mP2/3** | $1.0 \pm 0.3$ | $0.54 \pm 0.1$ | $5 \pm 2$ | $0.5 \pm 0.1$ |
| | **T/PK** | $21 \pm 2$ | $90 \pm 30$ | $1000 \pm 200$ | |
| | **ΔT/PK** | $2.7 \pm 0.7$ | $1.4 \pm 0.1$ | $16 \pm 7$ | $1.4 \pm 0.2$ |
| | **ΔP4** | $38 \pm 4$ | $35 \pm 6$ | $560 \pm 70$ | |
| | **ΔP6** | $16 \pm 6$ | $8.1 \pm 0.7$ | $130 \pm 30$ | |
| | **P4/5/6** | $6 \pm 3$ | $3.0 \pm 0.2$ | $34 \pm 9$ | $4 \pm 0.2$ |
| | **ΔU** | $12 \pm 2$ | $2.4 \pm 0.3$ | $70 \pm 10$ | |
| **LaM** | **1-268** | $130 \pm 10$ | $41 \pm 3$ | $1200 \pm 300$ | $70 \pm 30$ |
| | **mP2/3** | $12 \pm 3$ | $9.4 \pm 0.3$ | $70 \pm 40$ | $2.6 \pm 0.3$ |
| | **T/PK** | $5 \pm 1$ | $10 \pm 1$ | $170 \pm 20$ | |
| | **ΔT/PK** | $340 \pm 70$ | $140 \pm 30$ | $2000 \pm 500$ | |
| | **ΔP4** | $170 \pm 40$ | $80 \pm 10$ | $2100 \pm 300$ | |
|  | **ΔP6** | 13 ± 3 | 5.6 ± 0.9 | 30 ± 20 | 3.1 ± 0.7 |
|  | **P4/5/6** | 130 ± 30 | 8 ± 2 | 600 ± 100 | 18 ± 4 |
|  | **ΔU** | 4 ± 1 | 5.6 ± 0.8 | 20 ± 10 | 2.6 ± 0.4 |
| **xRRM** | **1-268** | 28 ± 3 | 6 ± 1 | 220 ± 30 |  |
|  | **mP2/3** | 3 ± 1 | 2.6 ± 0.5 | 19 ± 8 | 1.0 ± 0.1 |
|  | **T/PK** | 18 ± 5 | 30 ± 5 | 590 ± 90 |  |
|  | **ΔT/PK** | 4 ± 1 | 2.3 ± 0.2 | 28 ± 5 | 1.4 ± 0.6 |
|  | **ΔP4** | 4 ± 1 | 2.4 ± 0.3 | 40 ± 10 | 2.1 ± 0.8 |
|  | **ΔP6** | 4 ± 1 | 1.4 ± 0.2 | 14 ± 7 | 1 ± 0.7 |
|  | **P4/5/6** | 18 ± 5 | 7.6 ± 0.9 | 90 ± 30 | 8 ± 1 |
|  | **ΔU** | 4 ± 1 | 1.4 ± 0.3 | 20 ± 10 | 0.18 ± 0.004 |
| **F331A** | **1-268** | 8 ± 2 | 1.8 ± 0.3 | 24 ± 3 | 3 ± 1 |
| **D333R** | **1-268** | 3 ± 1 | 1.2 ± 0.1 | 29 ± 4 | 0.5 ± 0.1 |
| **R344A** | **1-268** | 48 ± 7 | 14 ± 4 | 550 ± 60 |  |
| **W387A** | **1-268** | 40 ± 4 | 2.7 ± 0.9 | 230 ± 50 | 4 ± 1 |

According to MST experiments, AtLa1 binding to AtTR is comparable in affinity to the minimal AtTERT RNA binding domain TRBD (Figure S4C, S4E), and to other pol-III transcripts, with these and other affinities summarised in Table S3. To explore this further, *in vitro* experiments were performed to attempt to map the location of AtLa1-AtTR interaction at a molecular level.

### La1 binds to telomerase RNA *via* its C-terminal xRRM domain, stabilised by LaM

Like many genuine La proteins and most LARP7 proteins, AtLa1 has a nonstandard RRM near its C-terminus, in addition to the La motif and adjacent RRM, which together form a La module (LaM) found in the majority of La and LARP proteins (Figure 1A). Sequence alignments with representative La and LARP7 proteins and Alphafold3 structural predictions indicate that this domain shares characteristic features of the extended helical RRM (xRRM) of *Tetrahymena* p65, including a RNP3-like RNA recognition sequence, a lack of single-stranded RNA binding sequences RNP1 and RNP2 found in canonical RRMs (Figure S5A, (31)) and a third predicted alpha helix in addition to the two found in canonical RRMs (Figure S5A). Genuine La proteins, except recently reported *in vitro* experiments for human LARP3 (9), have not been implicated as having a specific role in binding TR, however the LARP7 family, absent in plants, has evolutionarily diverse members with well-characterised TR binding properties (2). AtLa1, has a similar predicted domain structure (Figure1A) to TR-binding *Schizosaccharomyces pombe* Pof8 (30,31) and *Tetrahymena thermophila* p65 (59). Small angle X-ray scattering (SAXS) data are consistent with these predictions, and correlate to an ensemble of models where structured LaM and xRRM domains are freely mobile in solution, joined with a flexible linker (Figure S1D, Figure 8A,B, Table S6).To determine which domain(s) in AtLa1 are responsible for AtTR binding, the protein was expressed as a selection of fragments containing one or more of these domains (Figure 1A, Figure S1A, S1B). EMSAs detect a change in apparent AtTR mass consistent with complex formation in fragments containing the C-terminal amino acids 236-433, including the minimal fragment xRRM itself, which exhibits a modest, but distinct shift consistent with its lower mass compared to full-length (FL) or 106-433 fragments (Figure 2B). In contrast, fragments lacking these residues show no change in apparent AtTR mass over the concentration range explored. For a quantitative measurement of AtTR binding, MST was employed (*K*_D_ values summarised in Tables 1 and S3), focusing on amino acids 1-206 which encompass the LaM and amino acids 236-433, which contain the xRRM (Figure 2C). Initial fluorescence measurements with AtLaM reveal 26x and 21x lower affinity for AtTR in the fast and slow processes respectively, compared to the FL protein, the difference in the T-jump fast process is even more pronounced for LaM having 60x less affinity for AtTR than the FL protein. Conversely, xRRM initial fluorescence has only 5x or 3x lower affinity for AtTR in the fast and slow processes respectively, and an 11x lower affinity fast T-jump process. This suggests that the xRRM has the bigger impact on AtTR affinity of the two domains, consistent with observations from EMSAs. For both fragments, the addition of 100x excess unlabelled budding yeast tRNA competitor has no effect on either initial fluorescence (beyond a lower range of absolute values, see supplementary information and Figure S4F) or T-jump data (Figure 2C, lower panel, dotted lines) suggesting that these isolated RNA binding domains have greater specificity than the FL protein. It is perhaps also noteworthy that the fast T-jump processes for AtLa1 FL and xRRM in the presence of competitor RNA are approximately the same, suggesting that the xRRM is at least responsible for the majority of specific AtTR affinity.

**Figure 2:**
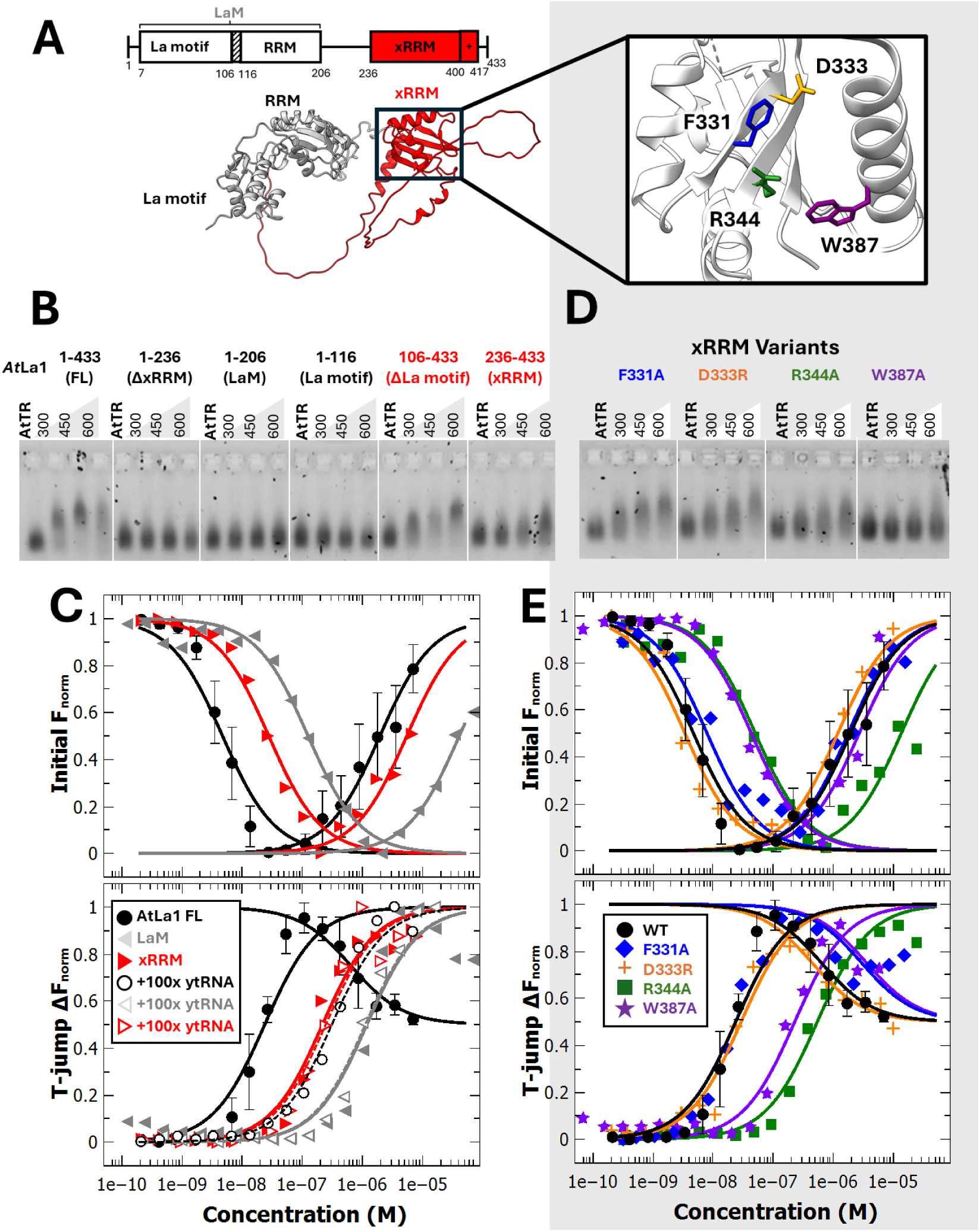
AtLa1 xRRM domain is predominantly responsible for AtTR binding. (**A**) Domain structure of AtLa1, including LaM (grey) and xRRM (red), as predicted by Alphafold2 (61), and visualized using ChimeraX (62). xRRM residues chosen for mutagenesis studies are shown in a magnified panel (the same colour scheme is used in B-E). (**B-E**) Binding of AtLa1 fragments (B, C) and AtLa1 xRRM point variants (D, E) to AtTR investigated by EMSA (B, D) and MST (C, E). (B, D) Representative band shift of 80 nM AtTR 1-268 caused by binding of AtLa1 protein constructs (concentrations in nM) is visualised in agarose gel by fluorescent staining. White lines have been added to separate panels in the same gel (full image in Figure S2). (C, E) Initial capillary fluorescence values from MST experiments (upper panels) or MST 1.5 s T-jump data (lower panels) for 3’ Cy5-labelled AtTR 1-268 and La1 fragments as indicated, with average AtLa1 FL data from Fig. 1F reproduced for comparison. Open symbols and dashed lines in (C), lower panel only, show data in the presence of 100x excess unlabelled tRNA competitor. Lines show equations fit to data used to calculate *K*_D_, error bars are the standard deviation.

The xRRM RNA-binding motif termed RNP3 found in human, alveolate and yeast xRRM domains has the sequence Y/W-X-D/Q (Figure S5A). The sequence in AtLa1 differs only in that the first amino acid is F, rather than Y or W. Given that both Pof8 and p65 also bind to their respective TR *via* their C-terminal xRRM domain (30,59), this suggests a conserved RNA binding mode shared by AtLa1 and LARP7 proteins. As such, we propose that the RNP3 motif (31) can include any aromatic residue, pending full structural characterisation. The xRRM fragment used here is partially disordered, but maintains a structured core with typical conserved features (Figure S1D, S8A,C, Table S6). Conserved AtLa1 xRRM residues F^331^, D^333^ and R^344^ are predicted to form part of beta-sheet 2 and 3 (β2, β3), and conserved alpha helix 3 (α3) residues Y^386^ and W^387^ are key parts of the p65 TR binding interface in *Tetrahymena* (26,33). Pof8 and p65 variants with RNP3 residues replaced had reduced binding affinity for TR (30,59). Structural models of these proteins also identify all these conserved residues present as part of the TR-binding side face of the xRRM (31,33) which is also present in models of AtLa1 predicted by Alphafold (60,61)(Figure S5B). Point mutant variants were prepared for F^331^, R^344^ and W^387^, where aromatic residues capable of pi-stacking interactions with nucleotides or phosphate backbone-attractive positively charged residues were replaced with non-binding A. Following the example of (30), D^333^ was instead replaced with R to create a D333R variant where this charge was reversed and not just annulled. EMSAs of AtLa1 point variants identified that R344A or W387A lower binding affinity clearly, whereas there is minimal change for RNP3 variants F331A or D333R (Figure 2D). MST experiments are consistent with this, in that F331A and D333R variants have initial fluorescence and T-jump data within error of wild-type (WT) AtLa1 (Table 1), whereas R344A and W387A show reduced binding (Figure 2E). For R344A and W387A initial fluorescence, the fast initial fluorescence process is impaired for both (9x or 8x respectively), whilst R344A also has lower affinity in the slow process (7x). In both cases, the fast T-jump process also has lower affinity, although for R344A it is 27x lower, compared to W387A where it is only 11x lower. This not only confirms that conserved xRRM residues R^344^ and W^387^ likely retain a structural role in TR binding in AtLa1, but also clarifies the technical point that differences in both initial PIFQ or PIFE processes are reflected in the *bona fide* binding detected by thermophoresis proper. The latter point is also consistent with the idea that fast PIFQ and slow PIFE processes arise from different molecular mechanisms.

### La1 binds to the stem-loop region between P5 and P6, and the three-way junction region of TR

Having determined the AtLa1 features responsible for AtTR binding, the next obvious question is which structural features of AtTR these RNA binding domains interact with. Of consideration were features typical for all TRs, such as the pseudoknot and template region (T/PK), specific AtTR features such as the P4/5/6 stem-loop and the three-way junction (TWJ) and regions typical for RNA Pol-III transcripts, such as 3’ poly-U tails (Figure 3A). Fragments and truncations of AtTR were synthesised *in vitro* and used for binding assays (Figures 3, S2, S3, S6, *K*_D_ values summarised in Tables 1, S2 and S3). Many of the AtTR constructs used here were previously used to demonstrate *in vivo* binding to AtTERT (21) giving confidence that they fold into biologically relevant structures. For several additional constructs, yeast-3-hybrid (Y3H) experiments under identical conditions as in (21) were performed with the TRBD (telomerase-RNA binding domain) constructs of AtTERT (Figure S3D), showing results consistent with the known model of AtTR-AtTERT binding (Figure S4G, (21)). As a first step in mapping the AtTR-AtLa1 interaction, competition ELISAs were performed with AtLa1 FL (Figure 3B, Table S2). These show most RNA constructs binding with similar preference as AtTR 1-268 (*K*_D_ = 41 nM), except for constructs with just the T/PK region (*K*_D_ = 190 nM) and just the P4/5/6 stem loop (*K*_D_ = 500 nM). This suggests that preferential AtTR binding may require the presence of the TWJ and/or 3’ and 5’-formed P1a stem that is absent in both constructs. Regarding the biological role of La proteins and the question of the 3’ terminus of AtTR, including any poly-U sequences previously suggested to be important for La1 binding, ELISA competition experiments were performed using AtLa1 FL and AtTR constructs synthesised with many variant ends (Figures 3G, S3B), including that first reported for ‘ATR8’, 1-259 (20) and those with or without poly-U motifs (Figure S3C). Affinities only vary by a factor of 10, (K_D_ = 9 nM to 90 nM) but seem to follow a pattern that relates to the length of the 3’ end or conversely the amount of the 5’ end that is exposed, considering that both termini form the closing P1a stem together assuming published models of TR (15) are accurate. The model for AtLa1 binding is therefore justified in including this P1a stem, with some amount of ssRNA overhang presumably affecting recognition or stability of interaction in this experiment, rationalised by the idea that RRM binding is sensitive to ssRNA content. EMSAs were also performed for all AtLa1 domain fragments and mutants (Figure S2), to compare AtLa1 1-268 and 1-262 missing the 3’ poly-U trailer (ΔU). Consistent with observations for pre-tsnoRNA, EMSAs with ΔU appear unchanged, suggesting that the short poly-U trailer is largely unimportant for AtLa1-AtTR binding. To map the site of AtTR binding in detail, AtTR fragment-binding MST experiments were performed with AtLa1 FL, followed by the LaM and xRRM fragments to examine the role of the individual domains. As previously, analysis will consider initial fast PIFQ and slow PIFE processes which so far seem to relate to specific protein-RNA interactions, together with the fast T-jump process which relates to *bona fide* binding.

**Figure 3.**
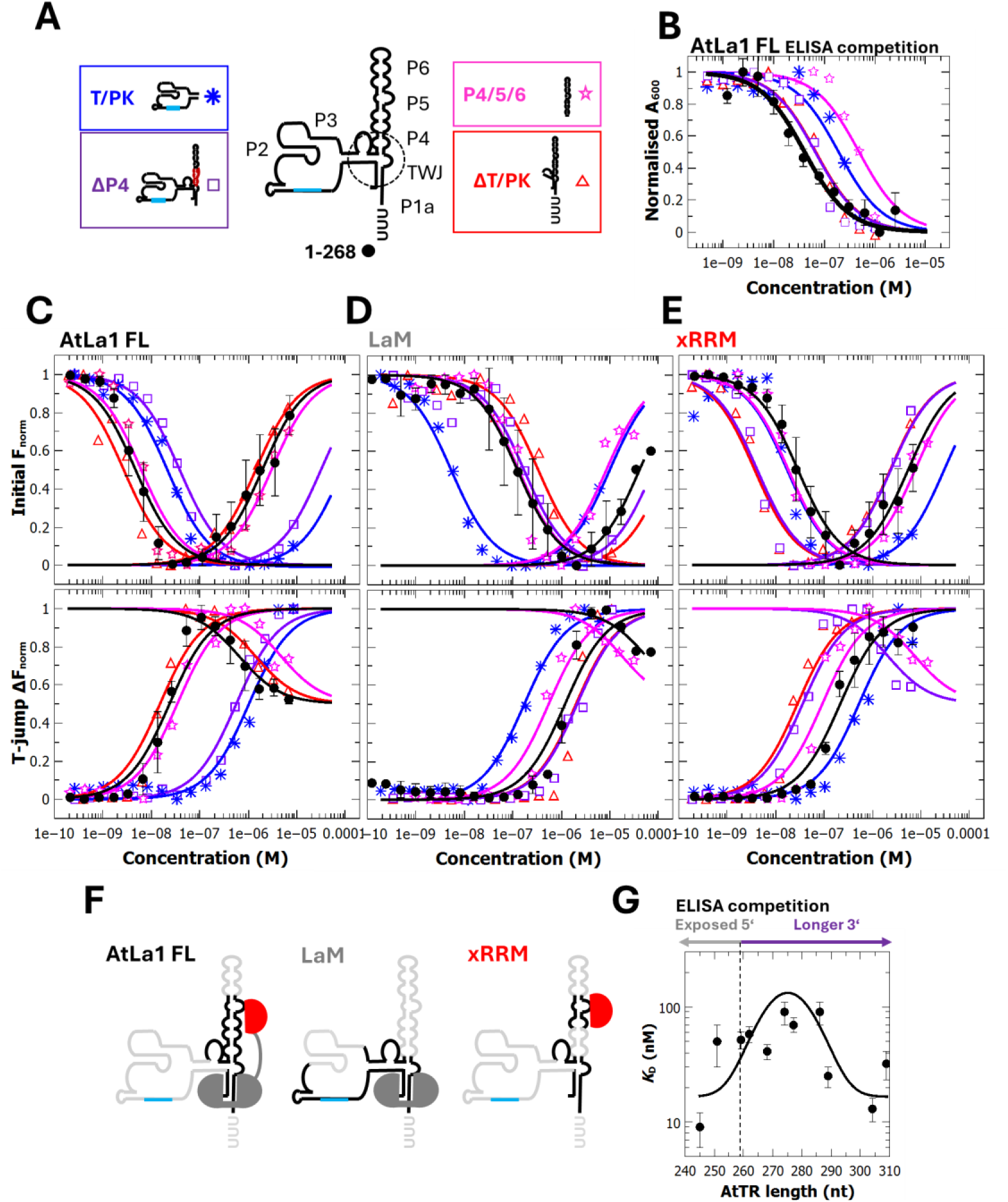
Mapping La1 domain binding to AtTR. (**A**) Overview and legend of representative AtTR constructs used in La1 binding experiments (B-E, additional constructs in Figure S6A), template (cyan) and deleted P4 (red) region are highlighted. (**B**) ELISA competition experiments detecting immunolabelled AtLa1 bound to ≤0.5 pmol biotinylated AtTR 1-268 and non-biotinylated competitor AtTR fragments. (**C, D, E**) Initial capillary fluorescence values from MST experiments (upper panel) or MST 1.5 s T-jump data (lower panel) for 5 nM 3’ Cy5-labelled AtTR fragments and La1 FL (C), LaM (D) or xRRM (E). Lines show equations fit to data used to calculate *K*_D_. (**F**) Schema of AtTR regions which bind AtLa1 LaM (grey) or xRRM (red) domains. Possible binding sites are shown in black, unlikely/nonspecific binding sites in light grey. (**G**) *K*_D_ values from AtLa1 ELISA competition experiments as (B) but with varying AtTR 3’ termini. The line shows fitting to a Gaussian distribution centred on 275 nt, full details in Figure S3B-C.

For AtLa1 FL, there is reduced affinity in all of these processes (initial fast PIFQ, initial slow PIFE and slow T-jump) for the T/PK construct, compared to 1-268 (Figure 3C, Table 1). This suggests that the T/PK is dispensable for FL binding, consistent with ELISA data, and with an observation that the construct ΔT/PK and the P4/5/6 stem have initial fluorescence and T-jump data comparable with AtTR 1-268. Similar reduced affinity is observed for a construct missing the P4 region and the poly-U tail (ΔP4, (21)) suggesting that P4 is part of or close to a binding site. A construct shortened by the distal part of the P6 region (ΔP6, Figure S3A, Table S3) also has lowered affinity in all MST characteristic processes, consistent with the idea that there is a binding site on the P4/5/6. Paired regions of P4/5/6 are formed by guanine doublet/triplets (Figure S4G-H). Thus, we investigated AtLa1 binding to several AtTR variants mutated within these regions. Modification of AtTR guanines 185-186 to cytosine, reduces the affinity of the initial fluorescence processes 3x, leading to a 7x loss in fast T-jump affinity compared to AtTR 1-268 (Figure S4H, Table S3), whereas there is no change when other guanine stretches along the P4/5/6 stem are modified, consistent with the idea of a binding site between P5 and P6 stems. MST measurements (Figure S6) seem to offer greater resolution than EMSA data (Figures S3A, S2) regarding the AtTR ΔU construct compared to AtTR 1-268, which shows a modest loss of affinity for AtLa1 FL (Figure S6B, Table S3). This suggests that the role of the short 3’polyU trailer in AtLa1 binding cannot be entirely discounted, but at most has a relatively subtle effect compared to other parts of TR. Finally, a construct modifying the P2 and P3 base pairing regions so that the pseudoknot cannot form (mP2/3, Figure S6) was investigated. The mP2/3 construct is catalytically inactive (15) and unable to bind TRBD (21), however, it has slightly higher affinity fast and slow initial fluorescence processes (5x and 3x, respectively) and a higher affinity fast T-jump process (3x), compared to AtTR 1-268. This suggests that disrupting the highly structured pseudoknot region may allow the formation of alternative structures that might be a possible artefact target for RNA-binding proteins.

The most striking difference between AtLa1 FL binding to various AtTR constructs and binding of LaM is its preference to bind the T/PK construct (Figure 3D) as well as ΔP6, ΔU and mP2/3 (Figure S6C) constructs (compare also Table S3). Removing the possibility of the pseudoknot structure formation in the mP2/3 does not affect LaM binding. This suggests that overlapping parts among these constructs might be involved in specific recognition by this protein domain. Namely, TWJ-adjacent regions of T/PK and possibly the P1a stem (Figure 3F). The binding affinities of xRRM to AtTR constructs are similar to AtLa1 FL, and quite opposite to LaM. Only T/PK has reduced binding affinity compared to AtTR 1-268, having a 5x lower affinity slow initial fluorescence process and a 2x lower fast T-jump process (Figure 3E). All other constructs bind with 1-268-like affinity or greater, suggesting that overall T/PK, P6, 3’polyU and curiously even the P4 stem are dispensable for xRRM binding (Figure S6D, Table S3).

Considering these observations together, it is clear, especially from ELISA competition experiments that AtLa1 FL likely has two binding sites on AtTR, consistent with having two structured RNA-binding domains (Figure 3F). The FL protein does not require either the distal P6 end of the P4/5/6 stem or the T/PK for high-affinity binding. The LaM likely recognises the adjacent regions of TWJ, perhaps together with the templating region and the connecting stem-loop and P1a stem. The xRRM binding is sensitive to the central region of the P4/5/6 stem-loop between P5 and P6, although it cannot be ruled out that it has some interaction with the TWJ adjacent regions. Assuming that the domains both sterically hinder each other from binding to identical or adjacent sites, this therefore suggests a model where one domain binds to the mid-P4/5/6 stem, between P5 and P6 (probably xRRM) and another near to the TWJ (probably LaM), perhaps stabilised by the adjacent stem-loop leading to the T/PK. The observation that the ΔP4 construct of AtTR binds poorly to AtLa1 FL, but is not involved in LaM or xRRM binding, is also rationalised if the absence of this stem merely brings the binding regions close enough that there is steric hindrance for both binding events to take place.

Overall, this fragment analysis suggests that AtLa1 binding to AtTR very much resembles the binding of *Tetrahymena* p65 to its equivalent TR. Given the similar affinity for AtLa1 and diverse plant TRs, this model of La binding to TR may also apply to many, if not all land plants. Another similarity between ciliates and plants is the presence of DUF3223 found in AtDomino, the one known protein interactor of AtLa1 (12).

### Domino contains a novel RNA-binding domain and binds to a three amino-acid motif in the La1 RRM

AtLa1 was previously reported to bind to AtDomino, a plant-specific protein of unknown function, which also binds AtTERT *via* its telomerase-specific N-terminal domain (12). Given that AtDomino shares an otherwise unique RNA-binding domain with the genuine La proteins of ciliates, and currently it is the best-characterised protein interactor of AtLa1 (12,35), experiments were performed to further investigate the molecular interface between these proteins.

To initially identify the domain(s) in AtLa1 responsible for AtDomino binding, the yeast 2-hybrid (Y2H) assay was used with the same library of AtLa1 fragments generated to study AtLa1-AtTR binding (Figure 4A). Consistent with previous reports using this technique (12), AtLa1 FL binds AtDomino strongly, as evidenced by growth in the presence of up to 30 mM aminotriazole, compared to an empty AD vector control (Figure S7A). Similar binding was observed for AtLa1 fragments LaM (aa 1-206), ΔxRRM (aa 1-236) and ΔLa motif (aa 106-433). In contrast, fragments La motif (aa 1-116) and xRRM (aa 236-433) show much reduced growth, suggesting that residues 116-206 contain the critical region for AtDomino binding. This is also the region of the AtLa1 sequence, which forms the canonical RRM domain, and Alphafold3 (60) predictions of the AtLa1-AtDomino complex model a loop containing residues S^124^, Y^125^ and D^126^ as the main binding interface (Figure 4A). These residues were replaced with glycine in the La1 variant SYD124GGG, and the binding of AtDomino was investigated using both *in vivo* Y2H assays and quantitative *in vitro* ELISA. The results from Y2H assays show that La1 SYD124GGG is no longer capable of binding AtDomino (Figure 4B), further confirmed with semi-quantitative Y2H beta-galactosidase assays (Figure 4C). ELISA experiments (Figure 4D) demonstrated a more than 10-fold reduced affinity for AtDomino in the SYD124GGG variant, compared to WT AtLa1. It is therefore assumed that the Alphafold3 prediction of the AtLa1-AtDomino structure is accurate in this regard. In contrast, AtLa1 xRRM variants with reduced AtTR affinity or specificity (eg. R344A Figure 4B, others Figure S7A), bind AtDomino the same as AtLa1 WT, consistent with the prediction that AtTR binding and AtDomino binding both depend on different (domain) interfaces of AtLa1 (Figures 4E). AtDomino contains the same domain of unknown function present in the recently identified genuine La proteins of ciliates (2), which is proposed to be a novel RNA-binding domain. MST experiments with AtDomino and fluorescent-labelled AtTR reveal a change of fluorescence and thermophoresis similar to that observed for AtLa1 (Figure 4F), giving confidence that AtDomino is indeed an RNA binding protein. All MST characteristic processes for AtDomino show lower affinity for AtTR 1-268 compared to AtLa1. In the presence of 1000x excess unlabelled budding yeast tRNA competitor, the initial fluorescence processes slightly increase in affinity, although with the increase in noise and general higher error in MST data, it is difficult to think of this observation as being significant. In contrast, the fast T-jump process shows 4x lower affinity in the presence of competitor RNA. The AtDomino sequence has a large number of positively charged residues at the C-terminus (Figure 4E); to eliminate the idea that Domino binding is due to simple electrostatic attraction (eg. (63)) and not the presence of the domain of unknown function DUF3223, MST experiments were also performed with an AtDomino construct missing the C-terminus (DelC). DelC binds AtTR 1-268 with 4x lower affinity reported by the initial fluorescence change, but the slow process and T-jump data are otherwise within error of AtDomino (Figure 4F). As the N-terminal sequence of AtDomino is mostly residues expected to have RNA-repulsive negative charge at neutral pH, it is therefore likely that the RNA-binding properties of AtDomino predominantly arise from DUF3223. Repeats of key Y2H experiments confirmed that AtDominoDelC still comparably binds AtLa1 (Figure S7B), suggesting that DUF3223 is also the site responsible for protein-protein interactions with AtLa1 (Figure 4E), also consistent with structural predictions (Figure 4A). Initial MST experiments with AtDomino in the presence of AtTR fragments used to map AtLa1 binding do not show the clear-cut differences as observed for AtLa1, however (Figure S7D).

**Figure 4:**
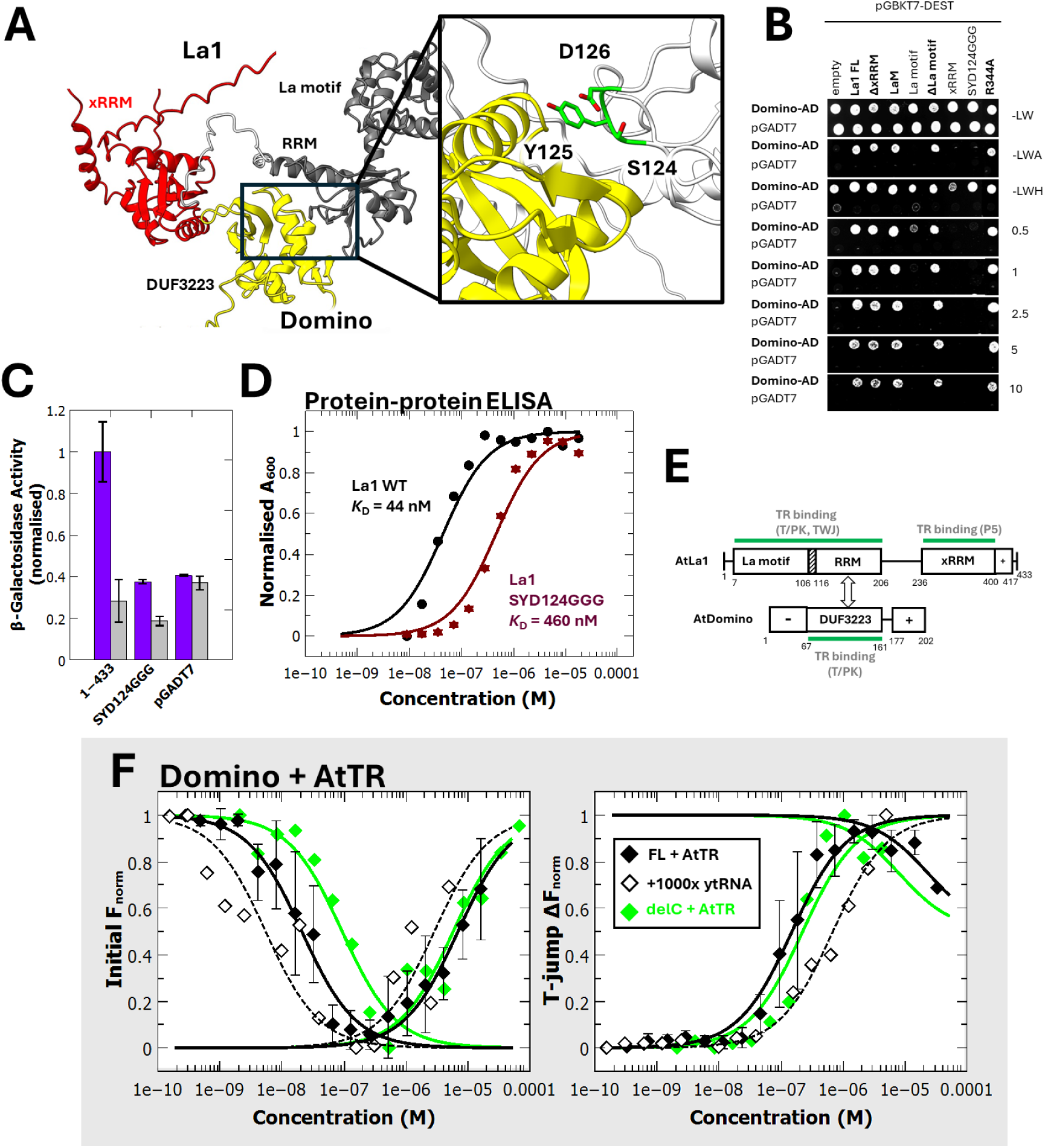
AtLa1 binds AtDomino *via* a three amino acid motif. (**A**) Alphafold3 prediction (60) of AtLa1-AtDomino complex showing AtLa1 LaM (La motif and RRM, grey) and xRRM (red), and AtDomino DUF3223 (yellow) domains. Highlighted panel shows interacting AtLa1 residues (green) on the RRM-DUF3223 interface. Structural visualisations prepared using ChimeraX (62). (**B**) Yeast 2-hybrid and (**C**) semi-quantitative beta galactosidase assays pairing Gal4 binding domain (BD) fused AtLa1 fragments and variants with Gal4 activation domain (AD) fused AtDomino. For (B), numbers refer to mM concentrations of 3-aminotriazole in the growth media, letters refer to amino acids absent for growth. For (C), BD-fused AtLa1 (purple bars) are compared to empty BD (grey bars), error bars represent standard deviation. (**D**) *In vitro* ELISA detecting AtDomino bound to immobilised AtLa1 WT (black circles) and AtLa1 SYD124GGG (dark red stars). Lines show equations fit to data used to calculate *K*_D_. (**E**) Domain structure of AtLa1 and AtDomino showing protein-protein (black arrows) and protein-RNA interactions (green bars), predicted domains as annotated, ‘+’ or ‘-‘ indicates regions of the sequence with correspondingly charged residues at neutral pH. (**F**) Initial capillary fluorescence values (left) and MST 1.5 s T-jump data (right) for AtDomino (black diamonds) and AtDomino delC (green diamonds) with 5 nM 3’ Cy5-labelled AtTR 1-268. For comparison, AtDomino interaction with Cy5-AtTR in the presence of excess unlabelled ytRNA competitor (open black diamonds, dashed lines) is shown. Lines show equations fit to data used to calculate *K*_D_, error bars represent standard deviation.

AtDomino therefore has both protein binding and RNA binding capabilities, which seemingly result from its single predicted structured domain. Although AtDomino binds to a semi-conserved loop in AtLa1 and has known interactions with AtTERT and other telomerase-related proteins (12), it still remains the question whether these binding activities have relevance to AtTR biogenesis.

### La1 enhances TERT binding to TR and forms a TR-binding complex with Domino

The next point to consider is whether AtLa1 binding to AtTR is compatible with AtTERT binding, which would explain why it is detected in AtTERT pulldowns (11). This question was initially addressed *in vitro* using MST in a three-component system (TRBD, AtTR, AtLa1), using 6xHis-tagged AtLa1 to eliminate the possibility of GST dimerization. In the presence of 125 nM (Figure 5A) or 1250 nM AtLa1 (Table S4), the minimal AtTERT telomerase RNA binding domain (TRBD, (21) has a considerable increase in affinity to AtTR in the slow initial fluorescence change (28x or 41x, respectively). Curiously, the fast fluorescence change and fast T-jump processes are not detected over the concentration range of the experiment. These data could indicate that interactions are complete at the minimum protein concentration; the lack of T-jump is also rationalised if there is no further change in mobility of the complex in solution in the presence of both AtLa1 and TRBD, such as if one protein binding instantly displaces the other, or the mobility of the trimolecular complex improves due to conformational change.

**Figure 5:**
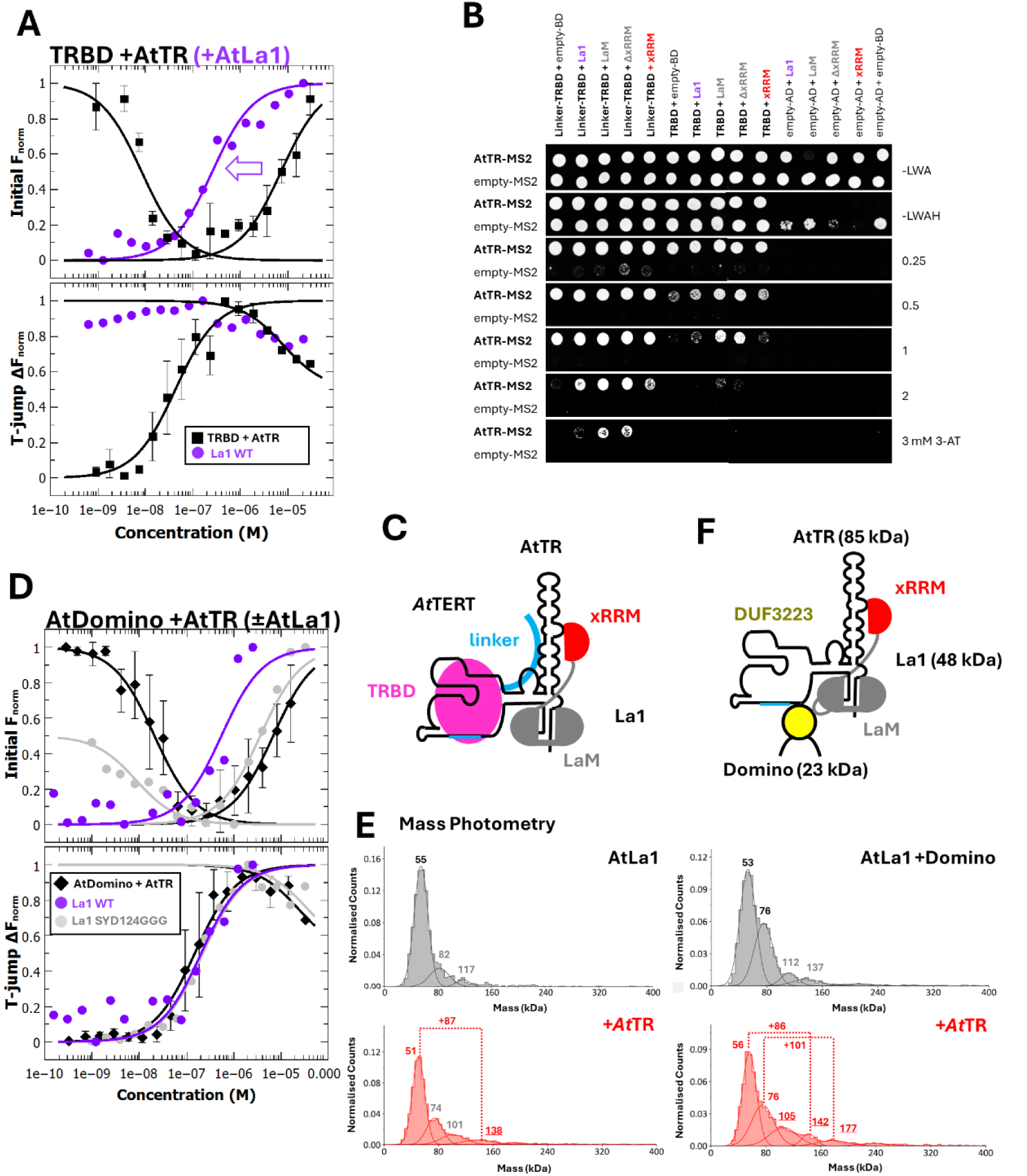
RNA binding complexes of AtLa1. (**A,D**) Initial capillary fluorescence values from MST experiments for 5 nM 3’ Cy5-labelled AtTR 1-268 and (A) AtTERT minimal binding domain (TRBD) or (D) AtDomino, with or without constant AtLa1 as annotated. AtDomino average data are reproduced from Fig. 4F, TRBD average data are reproduced from Figure S4E. Lines show fittings used to calculate K_D_ values, error bars represent standard deviation. (**B**) Multicomponent yeast three-hybrid experiments pairing AtTR 1-268 MS2 or MS2 only with AD constructs of AtTERT TRBD variants in the presence of AtLa1 BD or empty BD constructs, as annotated. Numbers refer to mM concentrations of 3-aminotriazole in growth media, letters refer to amino acids absent for growth. (**C,F**) Schemas of (C) partial AtTERT-TR-La1 complex omitting AtTERT TEN, RT and CTE domains, (F) possible AtTR-La1-Domino complex, with domains as annotated, LaM in grey, xRRM in red, DUF3223 in yellow, TRBD in pink and RNA-binding linker in cyan. (**E**) Mass photometry (MP) deconvoluted mass spectra of samples with AtLa1, AtLa1+AtDomino with or without AtTR, as annotated, Solid lines show Gaussian fittings with values in text, annotated with average mass values in kDa, numbers in grey are peaks which cannot be unambiguously assigned (more details in Table S5).

To investigate whether these *in vitro* observations can also occur *in vivo*, and to test our interpretation of MST data, experiments were then performed using a multi-component yeast 3-hybrid system (Figure 5B). These experiments are similar to those performed previously to investigate AtTERT-AtTR binding (21), with the addition of constructs encoding for additional proteins (taken from the Y2H system; Figure 4B) in the same cell. In this experiment, BD-fused constructs are not used to test the canonical AD-BD interaction as in Y2H, but rather serve as a means of selecting for the presence of the La1 protein in yeast cells (Figure S7F). For these experiments, TRBD was investigated with or without an unstructured linker that stabilises TR binding (21). Consistent with our interpretation of MST data, yeast with the AtTR-MS2 construct, TRBD AD constructs and AtLa1 BD can tolerate growth in the presence of higher 3-aminotriazole (3-AT), compared to when an empty BD replaces AtLa1. Curiously, the effect is even more pronounced for yeast with AtLa1 fragment LaM (aa 1-206) and LaΔxRRM (aa 1-236), suggesting that the LaM is the key domain for TRBD binding enhancement. This is consistent with the idea that a complex of AtLa1:TRBD:TR forms at least transiently, both *in vitro* and *in vivo* (Figure 5C), and that AtLa1 binding to AtTR serves to increase the probability of AtTERT binding telomerase subsequently, reminiscent of the stepwise system in Ciliates (64).

Our next question is whether AtDomino, AtLa1 and AtTR have an obvious interrelationship. MST experiments with AtDomino and AtTR 1-268 were therefore repeated in the presence of AtLa1 with a 6xHis tag. The presence of 125 nM (Figure 5D) or 1250 nM AtLa1 (Table S2) removes the fast initial fluorescence process and increases the affinity of the slow initial fluorescence process, similar to the observation for TRBD-AtTR. T-jump data do not change in the presence or absence of AtLa1. This suggests, first of all, that AtLa1 and AtDomino binding to AtTR are not mutually exclusive, as there is no indication of inhibition. There may also be a modest enhancement of binding as monitored by fluorescence, although this is not reflected in T-jump data, suggesting that this point is unclear. To investigate if the AtLa1-AtDomino interaction contributes to AtDomino-AtTR binding, AtLa1 SYD124GGG (Figure 5D), which does not bind AtDomino well, was employed in similar MST experiments. The presence of SYD124GGG only partially eliminates the fast initial fluorescence process of AtDomino-AtTR interaction, but does not otherwise change the remaining MST characteristics. Again, as there is no indication of inhibition, this suggests that AtLa1 and AtDomino binding to AtTR occurs at different sites and is not dependent just on protein-protein binding. This raises the possibility that AtLa1-AtDomino-AtTR could form a complex in solution.

To investigate whether the AtLa1:Domino:TR complex exists in solution, mass photometry (MP) experiments were conducted (Figure 5E, Table S5). This technique calculates the mass of molecules in solution binding to a laser-illuminated glass surface. Detection is based on interference contrast between reflected light and light scattered by molecules which scales linearly with the mass ((65), reviewed in (66)). For this experiment, 6xHis constructs were used to eliminate the possibility of GST dimerization, at the expense of a slight decrease in purity. MP has some relevant technical limitations, there is an approximately 30 kDa cutoff for masses detectable by this technique making AtDomino (predicted 23 kDa) undetectable as a monomer, and calibration of mass is based on proteins and therefore less accurate for nucleic acids (Figure S7E). In addition, the technique typically relies on low concentrations of protein (typically nM), meaning that complexes are likely to be present in low abundance unless they are exceptionally stable. Nevertheless, experiments where AtTR are present (Figure 5E, upper panels) show higher mass species present than those when it is absent (Figure 5E, lower panels), suggesting that protein-AtTR complexes are detectable.

The deconvoluted MP spectrum of AtLa1 alone shows a peak centred on 55 kDa (Figure 5E, upper left panel); this species is consistently present in all AtLa1-containing measurements and decreases in intensity when binding partners are present. We assign this peak as the AtLa1 monomer (predicted 48 kDa), and the minor high mass species as impurities or dimeric states. When equimolar AtTR 1-268 is present (Figure 5E, lower panels), a new peak appears at +87 kDa greater mass than the AtLa1 monomer, consistent with a 1:1 AtLa1:TR complex. Peaks previously assigned as impurities or dimers are similar, although may now include contributions from unbound AtTR.

Experiments were next performed with AtLa1 in the presence of a 10-fold excess of AtDomino (Figure 5E, right panels). The MP histogram shows several peaks; the major species are the AtLa1 monomer and a species at 76 kDa, which is much more abundant than the similar mass peaks recorded in the absence of AtDomino, which we therefore assign as 1:1 AtLa1:Domino (predicted 71 kDa). There are two more minor higher mass peaks, which could be dimers of the major peaks or otherwise combined impurities from both proteins.

When AtLa1 is in the presence of equimolar AtDomino and AtTR, the deconvoluted MP histogram now has new higher mass peaks, in addition to those assigned previously as AtLa1 and AtLa1:Domino. Peaks at 105 kDa and 142 kDa are likely to have background contributions from the impurities/dimers noted previously, but are nevertheless consistent with AtDomino and AtLa1 plus AtTR (predicted 85 kDa) respectively. The remaining unassigned peak at 177 kDa is broad (σ = 21 kDa) and low in abundance, but appears in no other measurements, and is most reasonably described as a complex containing at least one molecule each of each species present, AtLa1, AtDomino and AtTR (predicted 156 kDa).

Therefore, data are consistent with the formation of a 1:1:1 AtLa1:Domino:TR complex in solution at least, which exists alongside partial complexes of each of its components.

## DISCUSSION

### La1 binds polymerase III transcripts, but is not specific only to poly-U trailers

Previous studies showed that AtLa1 binds to polymerase III transcripts *in vivo*, specifically precursors for tRNA^Met^ and tRNA-snoR43.1, which both feature a 3’ poly-U tail (10). Data from this study described *in vitro* binding in the 100 nM range for several more polymerase III transcripts, consistent with the typical role of La proteins in binding these transcripts for protective or regulatory reasons (2,4,5,37). Of the transcripts tested here, AtTR has affinity and specificity comparable to pre-tsnoR43.1. This suggests that AtLa1 has a role in the early steps of plant telomerase biogenesis, however, copurification of AtLa1 with AtTERT (11) raises more possibilities that will be discussed below. Consistent with a proposed sensitivity to 3’ tails (10), the AtLa1 protein is somewhat sensitive to this part of AtTR, something that is known to be a key marker of polymerase III transcript recognition and maturation in eukaryotes ((5,6,37), reviewed in (3)). Unexpectedly, the experiments in this study show no obvious correlation between binding efficiency and the uracil content of the 3’ terminus of either AtTR, dicistronic pre-tsnoR43.1 or MRP1, although this preference is marked for constructs of snoR43.1 transcript mimicking two processing pathways suggested in (34). This suggests that AtLa1 binding specificity relies on more structural feature(s) than initially assumed, of which 3’ trailers are only part. As with many other La proteins, including *Tetrahymena* p65 (33) and *Trypanosoma brucei* genuine La (5), it is still the classic La module found in most genuine La proteins (2) that is likely to be responsible for interacting with the 3’ terminus of RNA. Different 3’ ends have been reported for AtTR by (20) and (15) and whilst there is a 10-fold variability in *K*_D_, suggesting a link to 5’ or 3’ terminus exposure, there is no obvious single preferred form for AtLa1 binding among the constructs tested here. There is the possibility that AtTR exists *in vivo* as variant molecules in different stages of 3’ processing or as variant molecules that function in different biological processes, for example telomerase (14) or heat-shock responses (20). It is also known that other plant TRs exist *in vivo* as multiple conformers, only one of which has the templating region exposed (67), a situation also recently reported in humans (68). In this case, AtLa1 may act as a method of selecting one or more forms of AtTR which are suitable for a role in telomerase. This could be similar to how fission yeast LARP7 protein Pof8 selects correctly-folded telomerase RNA with the aid of the LSm2-8 complex (30) or like the proposed human situation where H2A and H2B promote ‘open’ conformers (68). Moreover, TR exists as gene paralogs in many plant species; if plant La proteins are part of the quality control mechanism of TR, this suggests they may also be one part of what allows for adaptive evolution of these molecules ((69), reviewed in (16)).

### La1 fills the role of the missing LARP7 protein in plant telomerase

Identification of AtLa1 as a major species present in AtTERT pulldown experiments (11) raises the question why this protein is so prevalent and what its telomerase-related role might be. The former is explained by experiments where AtLa1 binds with high affinity (*K*_D_ < 100 nM) to AtTR and TR from several evolutionarily distant plant species. AtTR would certainly be expected in AtTERT pulldown experiments, so the presence of AtLa1 is explained by binding to this copurified AtTR. The model of AtLa1 binding to AtTR presented here features strong binding *via* the AtLa1 xRRM domain to the stem-loops between P5 and P6. This area has multiple predicted single-stranded loops with a high proportion of conserved nucleotides (15). Variation of the double-stranded regions changes AtLa1 affinity little, other than the P5 stem, which has a modest effect on binding. As such, the single-stranded sequences are likely the primary site of binding, consistent with what is known about xRRMs (70,71). Other plant TRs show conformational variation both *in vivo* and *in vitro* (67), including in these stem-loop regions, so it could be speculated that these loops may provide some sort of barcode or recognition site for AtLa1 binding, if this protein does have a role in selecting specific TR conformers. Ciliate LARP7 proteins, exemplified by *Tetrahymena* p65, bind to their respective telomerase RNA *via* very similar structural interfaces (28,33). This suggests that AtLa1 and perhaps other plant La proteins bind TR for a larger proportion of their lifetime, fulfilling one of the roles attributed to LARP7 proteins, which are absent in plants (2). AtTERT is known to bind AtTR *via* the PK region and the P4 stem (21), and we observe that AtLa1 increases the affinity of the AtTERT binding domain for AtTR both *in vitro* and *in vivo*. It might be tempting, therefore, to speculate that AtLa1 could act as a telomerase accessory protein as part of the *in vivo* active telomerase complex, something that is well characterised for yeast and ciliate LARP7 proteins (30,33,59). So far, only AtNAP57 (aka. Dyskerin or CBF5) has been proposed as a telomerase accessory protein in plants based on similarity to the human model (41), although its typical H/ACA RNA binding motif is not present in AtTR and it was not detected in AtTERT pulldown experiments (11). *In vitro*, AtNAP57 binds to approximately the same part of AtTR as AtLa1 with similar affinity (41). This is rationalized if these proteins are involved in different steps of AtTR maturation. Indeed, if plant TR is pseudouridinylated by the H/ACA snoRNP complex, of which AtNAP57 is proposed to be the catalytic subunit (72), this could be one explanation for a high affinity, but low duration binding interaction as part of AtTR biogenesis. *A. thaliana* also has another genuine La protein, AtLa2, although currently all that is known about this is that most likely it cannot complement the RNA maturation functions in yeast and does not bind pre-tRNAs and pre-tsnoRNA transcripts in vivo (10). From the perspective of our study it can be pointed out that the AtLa2 equivalent of the AtLa1 xRRM TR-interacting residue R^344^ is W, suggesting that this protein would not bind AtTR competently. Moreover, both genes are expressed *in vivo*, but differ in their tissue-specific expression with AtLa1 mRNA levels highest in tissues composed of actively dividing, i.e. telomerase active, cells. This is in contrast to AtLa2, which is more abundant in differentiated cells and senescent organs (see (10) and RNAseq analysis at TraVA database (73)),

### Plant La xRRM binding of TR is more like ciliates than yeast

The main driving force for AtLa1 binding of AtTR is its xRRM domain, distinguished from classic RRM domains by its additional helix and a change in RNP motifs (31). Typical RRM structure would suggest that aromatic residues are important to form a clamp for pi-stacking interactions, which passes along an RNA chain until a specific sequence binds according to the environment of the pocket (70,71). Based on sequence conservation and AtTR binding studies of point variants, we initially assumed that at W^387^ and F^331^ together form this clamp in AtLa1. Variation of the former decreases the affinity of binding, whereas the latter has no obvious effect on the AtLa1-TR interaction. Although the effect of modifying F^331^ is unexpectedly negligible, this is similar to the change in TR affinity of a minimal p65 xRRM domain where the corresponding Y^407^ is modified to A (59), but in contrast to the approximately 14-fold loss of yeast TR affinity where Y^330^ is modified to A in Pof8 (30). Pof8 binding to fission yeast telomerase RNA is in contrast with the plant/ciliate system, in that both LaM and xRRM have recognition sites on the equivalent of the T/PK loop, and 3’ binding is achieved by protein-protein interaction with the LSm2-8 complex that binds at this site (30). Given that yeast TRs are typically much larger (>1200 nt) than plant or ciliate equivalents (∼200-300 nt), it is of no surprise that a LARP7 protein of similar size to that in plants/ciliates requires additional help to span and fold this massive molecule. Modification of W^387^ in AtLa1 seems to modify only the fast initial PIFQ process, which we propose is related to substrate ‘scanning’ through an RNA-binding clamp (see below). As such, this residue is likely to be important for the mechanics of RNA binding, more than specific substrate recognition.

In addition to RRM clamp residues, adjacent residue R^344^ appears to have a more essential, conserved role in AtTR binding, presumably using positive charge for electrostatic attraction or steering of the substrate. Consistent with this, both initial PIFQ and PIFE processes are depressed when this residue is replaced, and we propose it has a role in both substrate ‘scanning’ and specific binding. Modifying the corresponding p65 or Pof8 R to A results in a 6-fold or 5-fold loss of RNA binding affinity, respectively, comparable to the 8-fold loss of binding for AtLa1. Another nearby residue, D^333^ has no obvious effect on binding affinity if it is switched from negative to positive, unlike the 8-fold loss of TR affinity for the corresponding Pof8 variant. In p65, the equivalent of this residue interacts with the aromatic ring of guanine-121, part of the GU-bulge essential for p65 binding (33) (Figure S8D), suggesting that this residue may be important for sequence/structure recognition in ciliates, but not plants.

Although hardly surprising, Alphafold3 predictions of the AtLaM, AtxRRM and AtTERT (60) overlay p65 and TERT well in a published structural model of *Tetrahymena* telomerase ((33), Figure S8D). Alphafold3 predictions of RNA structure still have very low confidence in our experience, so with current technology, it is probably not worthwhile considering AI models of AtTR or the AtTR-AtLa1 complex. However, the overall structure of the *Tetrahymena* equivalent of TR is broadly similar to AtTR, in that it has two main ‘arms’ joined by a TWJ, one of which is a T/PK region and the other is the stem terminus element (STE)(33), equivalent to P4/5/6. Positioning of AtLa1/p65 domains on *Tetrahymena* TR is consistent with our *in vitro* mapping of the AtLa1-AtTR interface (Figures 5F, S8B-D). Furthermore, there is clear space for AtTERT to bind within the main T/PK loop, which is unsurprising given the high conservation between TERT structures from different organisms. It is, however, likely that the exact positioning of the xRRM differs in the At system or is otherwise dynamic, considering that the nearest key xRRM residues to flipped-out RNA bases in the overlaid At structure are F331 and D333, both of which have no known impact on AtLa1-AtTR binding, whereas W387 does, and is situated a little far from RNA to interact (Figure S8).

### DUF3223 is a novel RNA-binding domain found in plants and ciliates

AtLa1 has one well-characterised protein binding partner, the DUF3223-containing protein AtDomino (11,12). Identified in this work is a semi-conserved SYD loop motif in AtLa1 that binds AtDomino, adjacent to the AtLa1 classical RRM domain. Experiments presented here also reveal that AtDomino binds RNA with nanomolar-range affinity, even when the positively-charged C-terminus is removed, confirming that DUF3223 in AtDomino at least has RNA-binding activity as previously speculated (2,36). AtDomino has an affinity for AtTR, comparable to that of AtLa1. Although we did not investigate a wide range of potential RNA substrates and cannot rule out interactions with other systems, there is evidence to suggest that AtDomino can form an AtTR-binding complex *in vitro*. The biological relevance of this complex is currently unknown, however. So far, DUF3223 has been found only in plant-specific DCL proteins (36,40), polymerase IV and V (38,39), and ciliate genuine La proteins, such as Mlp1 (2,37). Solution NMR structural information is also available for a bacterial D3223 protein (PDB 2K0M). In polymerase V, DUF3223 (referred to as DeCL) is found in the largest subunit NRPE1 (38). It binds the 3’ exonuclease RRP6L1 and is essential for *in vivo* RNA-guided DNA methylation and gene silencing, but is dispensable for core activity *in vitro* (39). As processing of human TR precursor involves participation of RRP6 (74), this raises the question of whether AtLa proteins have a role in the activity of plant-specific proteases, or whether different DUF3223 domains carry structural specificity for select roles.

AtDomino, like other DCL proteins, is essential for plant survival beyond embryogenesis (36,40), and is proposed to have a role in ribosome biogenesis and cell division (36). AtLa1 is also essential for plant survival beyond embryogenesis, suggesting that both proteins may be involved in the same essential pathway (35). AtDomino has more protein-protein interactions with a variety of telomerase-related proteins, including AtTERT (12), suggesting a general role related to this system, perhaps in biogenesis of telomerase, a part of the active telomerase complex or as a flexible protein-RNA chaperone.

The *Tetrahymena* genuine La protein Mlp1 (Figure S5C) has a unique role in variant ciliate tRNA processing (37). This protein differs from classic La proteins, in that it does not have a traditional La module, instead it features only the La motif and a downstream DUF3223 (2,37). Sequence alignments with La and LARP7 proteins suggest a La motif adjacent xRRM domain is also present in Mlp1 in a region already known to bind RNA (37). Curiously, this proposed domain structure of Mlp1 (La motif, xRRM, DUF3223) is similar to the complex of AtLa1 and AtDomino. It could be speculated that AtLa1 has evolved to perform either LARP7-like functions or Mlp1-like functions depending on the binding of AtDomino, something that could be an interesting starting point for further studies.

### Microscale thermophoresis is a sensitive technique for investigating protein-RNA interactions

Inspired by previous protein-RNA binding studies using oligonucleotides (75) or non-specific long RNA binding protein domains (63), we used MST in this study to evaluate specific protein binding to structured non-coding RNA. One of our first observations was that we had biphasic changes in initial fluorescence, and biphasic processes in some of our thermophoresis data proper. The classic thermophoresis data are easily rationalised in terms of a transition between two species with different mobility in solution, such as that observed in the 1:2 binding stoichiometry in the model system of arrestin binding to DNA aptamers, which is well-described by macroscopic or microscopic binding equations (76). We also satisfied ourselves that initial fluorescence changes do not lead to artefactual differences in thermophoresis data as long as we stay within a range of fluorescence values already greater than the limits suggested by instrumental software (Figure S1F). Although aggregation, visible as traces deviating randomly from a predictable curve, is present in the majority of the MST traces we recorded for protein-RNA binding studies, this is rarely present for the initial few seconds of experiment time, including the 1.5 s T-jump. Equally, centrifugation of our samples essentially replicates thermophoresis results in many cases, suggesting that neither of our two biphasic processes result from a separate process of aggregation or precipitation. Crucially, the initial fluorescence processes are absent if the local-viscosity-sensitive Cy5 label is moved from the 3’ terminus to the 5’ (Figure S4E), suggesting that they genuinely report on structural changes in the RNA molecule. Having ruled out obvious artefactual origins for our results, we then considered the intrinsic photochemical properties of our chosen fluorophore, Cy5. Cyanine dyes already have well characterised protein-induced fluorescence intensity changes which have been exploited for other techniques (57,58). Consistent with this, we were able to replicate our MST initial fluorescence measurements in a conventional fluorimeter (Figure 1F, S1H). The slow initial fluorescence change can therefore be described as a classic PIFE process (58) where protein binding to RNA changes the structure of this molecule, leading to restriction of the Cy5 tag, and therefore enhancement of fluorescence. Consistent with this, circular dichroism can be used to track global RNA structural changes (Figure 1F, S1I), suggesting that PIFE in this case reports on a process largely free of non-specific contributions, compared to the T-jump.

In some cases, Hill analysis (Figure 1F, G) may be more appropriate for the fast processes, which may have cooperative binding behaviour. For now, we have elected to keep the fitting of binding equations as simple as possible in our analysis to allow for easy comparison of values, at the risk of oversimplifying things. Of the data presented here with 3’labelled RNA, the slow fluorescence PIFE process is only completely absent when other processes are also very low affinity (Figure S4B, R344A, W387A) suggesting that this process could always be theoretically detected given high enough protein to match affinity and an appropriately-placed Cy5 reporter fluorophore.

The fast initial fluorescence process is perhaps a little more difficult to rationalise, logically it is higher affinity than *bona fide* binding reported in thermophoresis, and represents a protein-induced fluorescence quenching (PIFQ, (57)). It decreases affinity modestly in the presence of competitor RNA, seemingly in a specificity-related manner (Fig. 1G). As such it may include non-specific contributions, but less so than thermophoresis. It also seems that the presence of one protein at sufficient concentration to ‘saturate’ RNA via this process (Figure 5A,D) can prevent its further detection in the presence of other RNA-binding proteins. This implies that all of the proteins tested here can perform this physical process interchangeably, albeit with different affinity and presumably different specificity.

PIFQ is described as arising from disruptions of fluorophore-labelled nucleic acid structure, thereby decreasing nucleic acid-induced fluorescent enhancement (57). For our protein-RNA system, there is a lack of circular dichroism change over the range of this PIFQ, so such structural changes to RNA would be local to the Cy5 tag and not disrupt the overall A-form structure. This could be explained if, prior to binding (and therefore higher affinity by definition), pseudocontact between molecules leads to changes in flexible ssRNA conformation (such as the 3’ trailer) that do not affect paired dsRNA helical regions or the general overall molecular shape. Although observed for non-RRM binding events as well, this is broadly consistent with the predicted mechanism of RRM domain binding to RNA (70,71), which features sequence-specific scanning of ssRNA (presumably responsible for PIFQ) prior to genuine binding based on sequence-related probability (giving rise to PIFE).

## CONCLUSIONS

Comparing the *in vitro* TR binding properties of AtLa1 with LARP7 proteins, in particular with Tetrahymena p65, there are consistent similarities with the protein domains and RNA interfaces involved. We found MST as a user-friendly and highly flexible technique for determining multiple properties of protein-RNA interactions, which are comparable to results from various other techniques. In particular, AtLa1 binding enhances AtTERT binding in a comparable way to the p65-initiated assembly of *Tetrahymena* telomerase. Based on these data, we hypothesise that a genuine La protein fills the role of otherwise-missing LARP7 in plants for the biogenesis of TR, in addition to its general role in RNA biogenesis. We cannot say yet whether La1 is part of the active telomerase complex, but our data so far are consistent with this possibility, pending further studies. Seemingly, of the few LARP7 proteins currently fully characterised there are examples of both those which are telomerase subunits, such as p65, p43 and Pof8 and those which are only involved in biogenesis, such as human LARP7. We also have evidence to support predictions that the DUF3223, present in the La-binding Domino protein, is a novel RNA-binding motif, together with its already-characterised protein binding capabilities. Currently, its exact role in the telomerase pathway, if any, is unknown, but we have identified a semi-conserved loop on plant La proteins that binds it, and the possibility of it binding AtTR alone, or in complex with AtLa1. Both the conservation of the role of a La protein in telomerase and the presence of DUF3223 seems to be a common trait of both plants and ciliates, hinting towards these both arising from common ancestral RNA-processing systems.

## DATA AVAILABILITY

All data are incorporated into the article and its online supplementary material. SAXS data are deposited to the SASBDB under the codes SASDYP2 and SASDYQ2.

## SUPPLEMENTARY DATA

Supplementary Data are available at NAR online.

## ACKNOWLEDGEMENTS

LJ is grateful to Jean-Luis Mergny and Václav Brázda (Institute of Biophysics CAS, Brno, Czech Republic) for advice on G4 characterisation and Jaroslav Malina (Institute of Biophysics CAS, Brno, Czech Republic) for help with spectroscopy. We would also like to acknowledge Jiří Fajkus (Masaryk University Brno, Czech Republic) for fruitful discussion and support. The CIISB, Instruct-CZ Centre of Instruct-ERIC EU consortium, funded by MEYS CR infrastructure project LM2023042 and European Regional Development Fund-Project „Innovation of Czech Infrastructure for Integrative Structural Biology“(No. CZ.02.01.01/00/23_015/0008175), is gratefully acknowledged for financial support for measurements at the CF Biomolecular Interactions and Crystallography. Special thanks go to Tomáš Klumpler and Josef Houser (Central European Institute of Technology, Brno, Czech Republic) for their advice, support and patience during LJ’s use of the core facilities. Molecular structures were aligned and visualised using UCSF ChimeraX, developed by the Resource for Biocomputing, Visualization, and Informatics at the University of California, San Francisco, with support from National Institutes of Health and the Office of Cyber Infrastructure and Computational Biology, National Institute of Allergy and Infectious Diseases.

## CRediT statement

Leon Jenner: Conceptualization, Investigation, Formal analysis, Methodology, Validation, Writing – original draft. Dzmitry Pruchkouski: Investigation, Methodology, Visualization. Writing – original draft. Barbora Stefanovie: Investigation, Validation, Visualization, Writing – original draft. Olga Novakova: Formal analysis, Resources. Monika Kubickova – Investigation, Validation, Formal Analysis. Petr Fajkus: Resources, Formal analysis. Marie Brazdova: Resources. Jan Palecek: Supervision, Resources, Writing – review & editing. Eva Sykorova: Conceptualization, Investigation, Resources, Supervision, Funding acquisition, Project administration, Writing – original draft, review & editing.

