## supplementary information for "La protein binding to telomerase RNA supports an evolutionary relationship between plant and ciliate telomerase pathways"

### Supplementary Data (Jenner et al. 2025)

#### (North)western blotting

Proteins were prepared and purified according to the description presented previously. Samples were resolved on a 12.5% SDS-PAGE gel, then transferred onto nitrocellulose membranes using a wet transfer system Miniprotean (Biorad) at constant current (350 mA) for 90 minutes. The transfer buffer consisted of 25 mM Tris, 192 mM glycine, and 20% methanol. Post-transfer, the membranes were stained with 0.1% Ponceau S (Carl Roth GmbH) for protein visualization, followed by destaining with deionized water until clear. To renature proteins, membranes were incubated in renaturation buffer (10 mM Tris-HCl, pH 7.5; 50 mM NaCl; 1 mM EDTA; 0.02% [w/v] Ficoll 400; 0.02% [w/v] polyvinylpyrrolidone-40; 1% [w/v] BSA) for one hour at room temperature with gentle shaking. The membranes were then washed three times with Northwestern (NW) buffer (renaturation buffer without BSA). Membranes were incubated in hybridization boxes with NW buffer containing $^{32}$P radiolabelled RNA and competitor RNA, where noted, either in vitro-synthesised control RNA or commercial tRNA from *Saccharomyces cerevisiae* (R5636, Sigma Aldrich). Incubation was carried out for two hours or overnight at room temperature with gentle shaking or rolling. Unbound RNA was then removed by washing the membranes 3–4 times with NW buffer for 10 minutes each. Following this, membranes were air-dried, wrapped in plastic foil, and exposed on an imaging desk (GE Healthcare). A Typhoon FLA 9000 phosphorimager (GE Healthcare) was then used to visualise bound $^{32}$P-labelled RNA.

For western blotting, the same procedure was followed until the Ponceau S destaining step. After this, membranes were blocked with 5% fat-free milk powder in 50 mM Tris-HCl, 150 mM NaCl, pH 8. They were incubated with 1:5000 diluted anti-GST primary antibodies (G1160, Sigma Aldrich) in the same buffer overnight, washed three times for 10 minutes with buffer only. Next, they were incubated with 1:8000 anti-mouse horseradish peroxidase fusion secondary antibodies (A0168, Sigma Aldrich) in the same buffer for 1 hour, washed three times as before and then visualised using a SuperSignal™ West Femto Maximum Sensitivity kit (#34096, Thermo Scientific) diluting the substrate 1:1 with water and then visualising chemiluminescence using an Amersham™ Imager 680 (GE Healthcare).

#### Fluorescence Spectroscopy

Fluorescence spectra of 10 µM N-methyl mesoporphyrin IX (Santa Cruz Biotechnology, sc-396879A), a kind gift from Václav Brázda (Institute of Biophysics CAS, Brno, Czech Republic), were recorded in 100 mM Tris-HCl, 100 mM KCl, pH 8, using a Varian Clary Eclipse fluorescence spectrophotometer (Agilent) and a 0.4 cm micro cuvette. RNA samples were added to NMM for a final concentration of 200 nM RNA, and fluorescence emission spectra were recorded with excitation at 378 nM, then plotted in QTIplot (IonDev Software), accounting for baseline drift and dilution. Fluorescence-monitored titrations instead used 20 nM AtTR-Cy5 in 50 mM Tris-HCl, 250 nM NaCl, 250 mM potassium acetate, pH 7 with 0.05% Tween, titrated with AtLa1 as indicated in figure annotations. Emission spectra were recorded using 625 nM as the excitation wavelength and 10 nm excitation and emission slit widths and then plotted as above.

#### Yeast protein expression analysis

Yeast cells were grown to OD~2.5 and lysed by incubation in 0.1 M NaOH for 5 min and boiling in SDS Laemmli buffer (62.5 mM Tris–HCl, 2% SDS, 5% β-mercaptoethanol, 10% glycerol, 0.002% bromophenol blue; (1)). Samples were separated by 12% SDS-PAGE, blotted, and analysed with antimyc-HRP (Abcam 62928) diluted to 1:5000 in 1% milk-TBST, and incubated for 1 hour at room temperature. After washing in TBST buffer, chemiluminescence was detected using SuperSignal™ West Dura Extended Duration Substrate (#34075, Thermo Scientific).

#### Small angle X-ray scattering

3 mg/mL AtLa1 or 1 mg/mL xRRM were purified with a 6x His tag to avoid contributions from globular fusion protein GST. SAXS data were collected using a Rigaku BioSAXS-2000 instrument at CEITEC (Brno, Czech Republic) equipped with a HyPix-3000 detector at a sample-detector distance of 0.48 m. Scattered intensity was measured in the range 0.009-0.65 1/Å, where $q = 4\pi \sin \theta/\lambda$; $2\theta$ is the scattering angle and $\lambda = 0.154$ nm. The datasets were normalized to the intensity of the transmitted beam and radially averaged using SAXSLab (Rigaku). Scattering curves from individual frames were checked for radiation damage and averaged. The corresponding scattering from the solvent-blank was subtracted to produce scattering profiles, which were then analysed using PRIMUS from the ATSAS 4.1:1-1 toolset (EMBL, Hamburg) to visualise data as a Kratky plot to evaluate protein folding. For further analysis, SAXS data were truncated from the first Guinier point to a maximum of q = 0.3 1/Å. EOM software (2) was used to fit theoretical scattering from ensembles of Alphafold-derived models to experimental data. The SAXS datasets and experimental details were deposited at SASBDB under the accession codes: SASDYP2 and SASDYQ2

#### Additional information for the evaluation of MST experiments

MST experiments were first performed with AtLa and a known *in vivo* dicistronic pre-tsnoR43.1 (Figure 1D), which is known to contain a 3’ UUU-OH trailer (3). Alternative substrates tested via MST (Figure S4A) were derived from putative pathways of pre-tsnoRNA processing proposed in (4) mimicking the processed transcript free of polyU trailer (pre-tsnoR43.1 ΔU), mature snoR43.1 ΔU variant, and putative intermediate product snoR43.1 retaining polyU trailer. AtLa1 and pre-tsnoR43.1 incubated together show comparable signals to AtLa1-AtTR with lower affinity in all MST characteristics, whereas affinity is about the same for a construct containing just snoR43.1 (Figure S4A). Constructs lacking the 3’ UUU-OH trailer (ΔU) had identical performance in the case of pre-tsnoR43.1 ΔU, but much reduced affinity in the case of snoR43.1 ΔU. AtLa1 had the poorest affinity for commercial yeast tRNA (ytRNA, used as a low affinity mature tRNA control) in all MST characteristics, thus the pattern of affinities is overall the same as that observed in ELISA competition experiments (Figure 1D). MST experiments where just the xRRM of AtLa1 (aa236-433) was in the presence of pre-tsnoR43.1 resulted in MST characteristics within error of those using full-length protein (Figure S4B), suggesting xRRM may be the primary binding site for this substrate. Consistent with this, when conserved xRRM residues $R^{344}$ or $W^{387}$ (Figure 2A, S5A) are changed to alanine, there is a decrease in affinity for all resolved MST characteristics.

Similar MST experiments were repeated with fragments of AtTERT, which are known to specifically bind AtTR (5). The minimal TR binding domain (TRBD, aa299-580) and the same fragment plus a linker that stabilises binding (linker-TRBD, aa229-580) have almost identical initial fluorescence and T-jump changes to AtLa1 and AtTR (Figure S4C, Table S3). When a truncated TRBD (TRBD-trunc, aa320-580) is employed, which was previously reported to abolish binding (5), all resolved MST characteristics have lower affinity. This gives confidence that both initial fluorescence changes and T-jump changes report on the strength of known protein-RNA interactions.

#### Sensitivity of the Cy5 fluorophore

To examine the specificity of AtLa1-AtTR binding, experiments were repeated in the presence of unlabelled competitor RNA (Figure 1G), either ytRNA or pre-tsnoR43.1, which, based on the results above, we would expect to be poor competitors or strong competitors, respectively. The presence of 1000x excess ytRNA does not alter the initial fluorescence changes a great deal after normalisation, whereas 10x pre-tsnoR43.1 does. However, in all cases and regardless of competitor, the range of initial absolute fluorescence measurements seems to decrease approximately according to:

$$\%\ decrease = \frac{ln(\frac{c_{comp}}{c_{labelled}})}{8}$$

Where $c_{comp}$ is the concentration of unlabelled competitor RNA, $C_{labelled}$ is the concentration of Cy5-labelled RNA, and % decrease is the percentage decrease in the total absolute signal range (Figure S4F). Attempts to instead fit these decreased data, normalised to values in the absence of competitor, are unconvincing, although perhaps due to the decreased signal range, data in the presence of competitors seem noisier.

Given the sensitivity that Cy5 dyes have to their environment and observations about the specificity of initial fluorescence changes, experiments were performed to compare 5'-labelled AtTR (3'TR, maleimide linkage to a sulfonated phosphate group) to the 3'-labelled AtTR used initially (3'TR, enzymatic addition of a labelled cytosine at the 3' terminus, Figure S4D,E). When 5'TR is incubated with AtLa1 (Figure S4D), there is now minimal change in initial fluorescence, only a decay of signal at high concentrations reminiscent of that observed for 3'TR and GST only (Figure S1K). T-jump data still show a typical biphasic profile; however, the fast and slow processes have lower affinity. In comparison, (linker-)TRBD incubated with 5'TR has a similar T-jump profile (Figure S4E). Despite this, there is no clear protein-dependent initial fluorescence change beyond a steady drop in signal at high protein concentrations, likely to be artefactual. This suggests that the position of the Cy5 tag in 3'TR, predicted to be at the end of a solvent-exposed ssRNA overhang (Figure S4G, (6), is both sensitive to structural changes but unlikely to interfere with AtLa1 or AtTERT binding. Conversely, having the Cy5 tag on the 5' terminus, predicted to be a single adenine adjacent to a dsRNA stem, is no longer sensitive to structural changes, perhaps due to the proximity of a rigid dsRNA region. In addition, Cy5 in 5'TR appears to inhibit AtLa1 binding, perhaps through steric hindrance if the P1a stem is important for AtLa1 substrate recognition. Consistent with this, AtTERT is known not to be sensitive to this region for binding, explaining its unimpaired binding to 5'TR.

Another observation was that high concentration samples of protein-AtTR have visible precipitation in solution, suggesting a change in solubility for the complex compared to where species are unbound. To explore this, AtLa1-AtTR (Figure S4D) and TRBD-AtTR (Figure S4E) MST experiments were performed using the same samples before and after centrifugation (5 min, 6000 g). The initial fluorescence values after centrifugation were in both cases lower than before, in a protein-dependent manner. The difference was plotted, normalised, and compared to the T-jump, obtaining very similar results, suggesting that differences in mobility due to sedimentation, thermophoresis, and electrophoresis can all detect complex formation comparably.

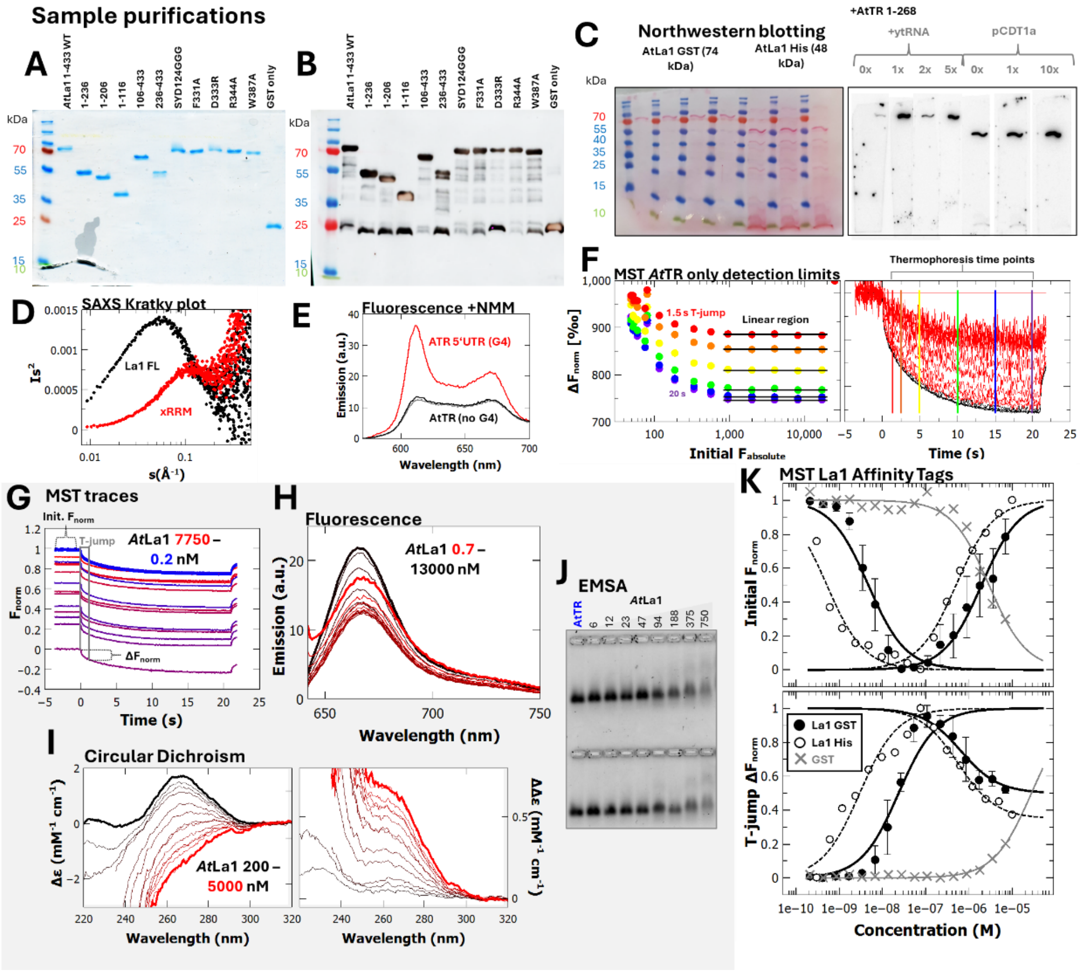

**<u>Figure S1</u> AtLa1 purification and AtLa1-TR binding extended data. (A)** Example Coomassie blue stained 12.5% SDS PAGE image showing purified GST-fused AtLa1 proteins used in this study. **(B)** Example anti-GST western blotting of a non-stained replicate of (A), 23 kDa species detected in most blots (but not typically in Coomassie gels of fresh protein) is assigned as traces of free GST. **(C)** Representative northwestern blotting of AtLa1 with C-terminal GST or 6x His, showing general Ponceau S staining of blotted proteins (left panel) and detection of radioactively labelled AtTR bound to AtLa1 (right panel) in the presence of non-labelled yeast tRNA (tRNA) or pCDT1a competitors with molar excesses as indicated. **(D)** Kratky plot of small angle X-ray scattering data of 6xHis tagged AtLa1 constructs as annotated (details in Table S6 and SASBDB under codes SASDYP2 and SASDYQ2) **(E)** Fluorescence emission spectra of 200 nM AtTR 1-268 (black) or ATR 5'UTR (red) RNA in the presence of 10 µM G4-sensitive dye N-methylmesoporphyrin (NMM), excited at 378 nM. Spectrum of 10 µM NMM alone (grey) shown for comparison. **(F)** MST detection limits of serially-diluted AtTR. (Left panel) thermophoresis of AtTR only, at a range of different absolute fluorescence values. Black lines show linear region where no artefactual ΔF is detected, with colours indicating time point of thermophoresis from 1.5 s (T-jump) to 20 s (i.e., 1.5, 2.5, 5, 10, 15, 20). (Right panel) raw data of the same experiment with coloured lines to indicate time points for analysis. Black MST traces are from the reliable linear dependence region, red traces show artefactual ΔF and noise where initial fluorescence is <1000 or lack of detectable thermophoresis where fluorescence is >250000. **(G)** Raw MST traces from representative AtLa1 to AtTR-Cy5 binding in premium coated capillaries to reduce aggregation, annotated to visualise calculation of initial fluorescence and thermophoresis values. **(H)** Fluorescence emission spectra of 20 nM AtTR-Cy5 excited at 625 nM and **(I)** Circular dichroism spectra of 50 nM AtTR-Cy5, titrated with AtLa1 as annotated. **(J)** Representative band shift of 40 nM RNA caused by binding of AtLa1 FL (concentrations in nM) as visualised in agarose gel by fluorescent staining. **(K)** (upper panel) initial fluorescence and (lower panel) T-jump values from MST experiments with AtLa1 tagged as annotated, or tag only, with Cy5-labelled AtTR 1-268, AtLa1 GST data are reproduced from Figure 1F for comparison. Lines show equations fit to data used to calculate $K_D$, error bars represent the standard deviation.

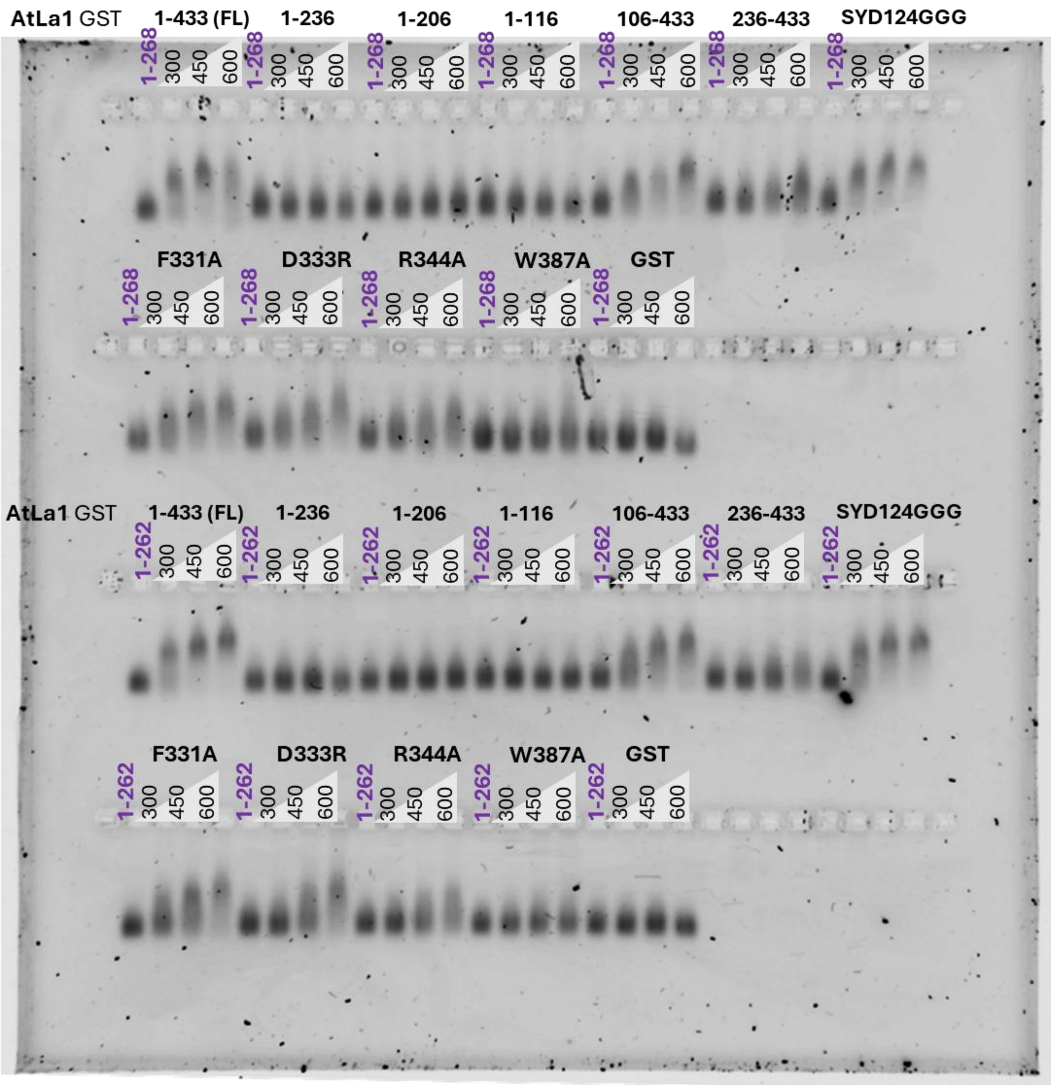

**<u>Figure S2</u> Full EMSA image of AtLa1 fragments and variants.** Representative EMSA using 1% agarose gels, 80 nM AtTR 1-268 (upper two rows), 1-262 (lower two rows), and AtLa1 fragments as annotated with concentrations in nM, visualised by fluorescent staining. Parts of this image are used in Figures 2B, 2D.

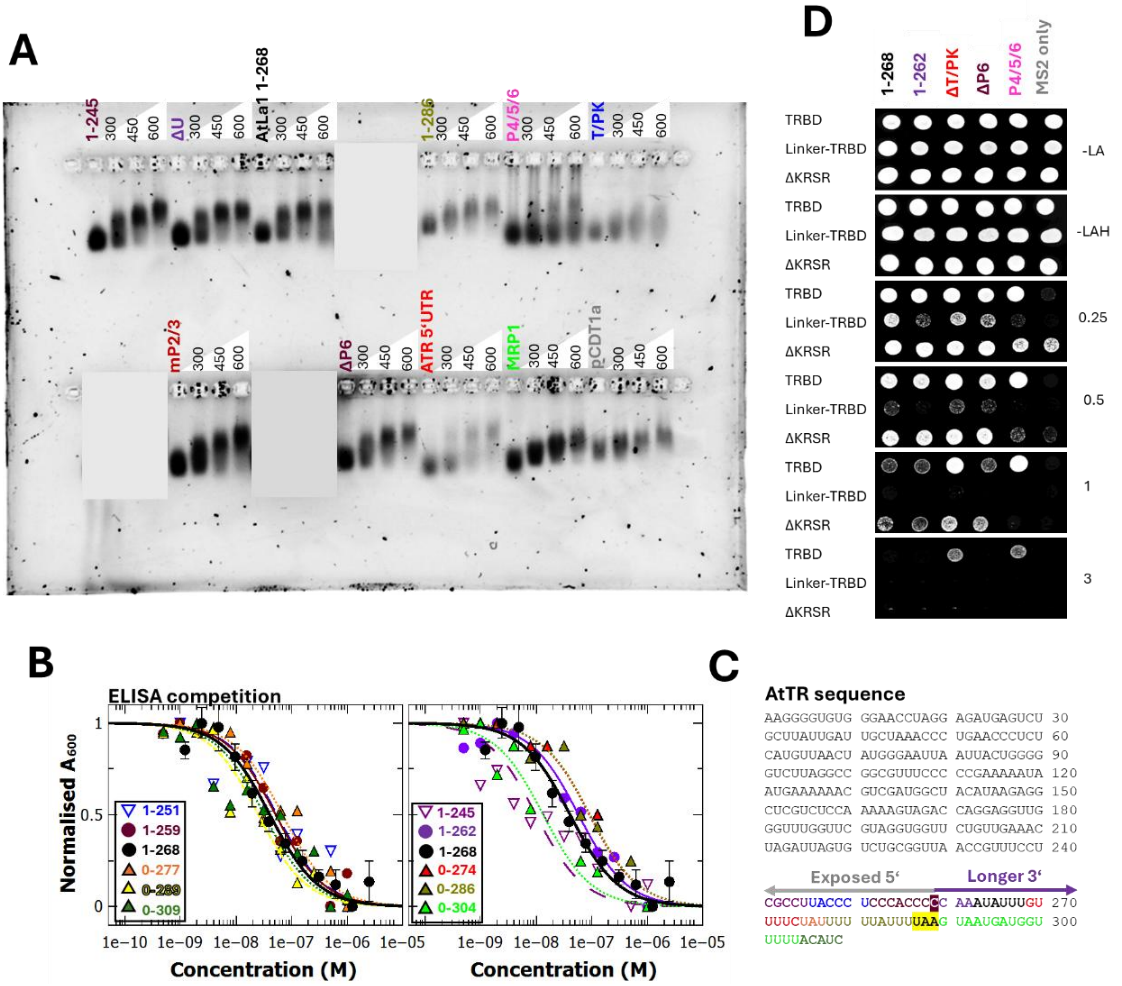

**<u>Figure S3</u> AtLa1 binding to variant AtTR constructs. (A)** Representative EMSA using 1% agarose gels, 80 nM AtTR constructs and other RNAs as annotated (160 nM T/PK, 320 nM P4/5/6 or ATR 5'UTR), incubated AtLa1 (concentrations marked in nM indicate where protein is added), visualised by fluorescent staining. Grey boxes obscure parts of this image not relevant to this work. **(B)** ELISA competition experiments detecting immunolabelled AtLa1 bound to ≤0.5 pmol biotinylated AtTR 1-268 and non-biotinylated competitor RNA as annotated, with data reproduced from Figure 1D for comparison. Lines show equations fit to data used to calculate $K_D$, error bars represent the standard deviation. For competition ELISAs, fittings were used to calculate $K_D$ values in Figure 3G. **(C)** Sequence of AtTR with colour coded endings as (B). Purple highlight shows cytosine 259 which is assumed to be the last duplex-binding base of the 3' terminus. **(D)** Yeast three-hybrid experiments pairing AtTERT constructs as annotated with MS2-tagged AtTR constructs as annotated, according to (5). Numbers refer to mM concentrations of 3-aminotriazole in growth media, letters refer to amino acids absent for growth.

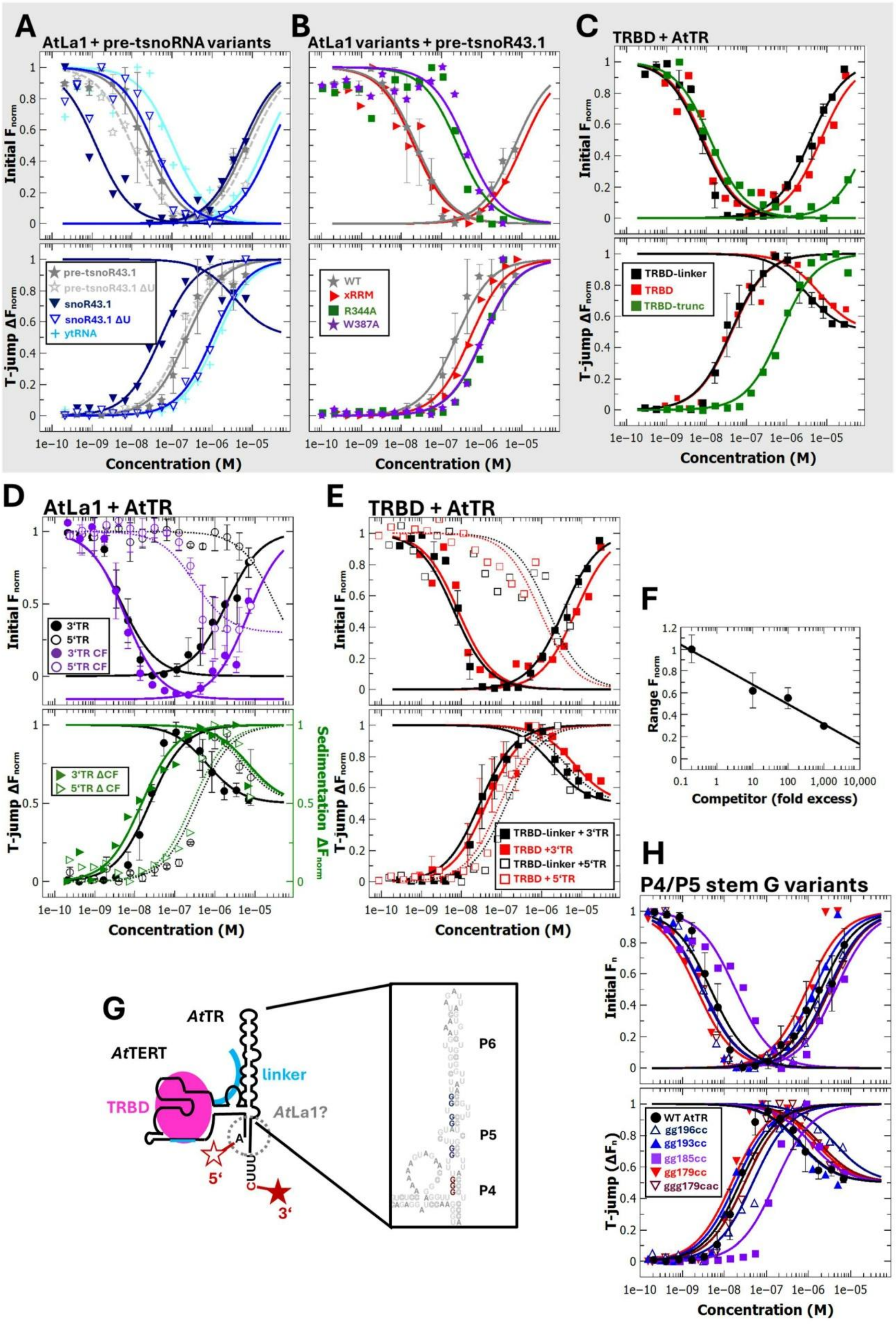

**<u>Figure S4</u> Microscale thermophoresis (MST) for protein-RNA binding experiments**. **(A-E)** (upper panel) initial fluorescence and (lower panel) T-jump values from MST experiments with **(A)** AtLa1 and Cy5-labelled pre-tRNA-snoR43.1 constructs as annotated, **(B)** AtLa1 variants as annotated with Cy5-

labelled pre-tRNA snoR43.1, **(C)** AtTERT TR binding fragments as annotated with Cy5-labelled AtTR 1-268, **(D-E)** AtLa1 FL (D) or the AtTERT TR binding domain (TRBD ± RNA-binding linker)(E), and 3' or 5' Cy5-labelled AtTR, before or after centrifugation (5 min, 6000 g) as annotated. TRBD-TR data from (A) are reproduced in (E), AtLa1-TR data from Figure 1F are reproduced in (D) for comparative purposes. Lines show equations fit to data used to calculate $K_D$, error bars represent the standard deviation. Data after centrifugation (D) are normalised to pre-centrifugation values and go below 0. **(F)** Logarithmic relationship between the normalised range of initial fluorescence values and fold excess of unlabelled competitor in MST experiments. **(G)** Schema of partial AtTERT binding to AtTR, showing alternative positions of Cy5 fluorophores (red stars) as annotated and possible region of 5'Cy5 inhibition of AtLa1 binding (grey dashed circle). A box highlighted a 2-dimensional structure of the AtTR modelled after that proposed in (6) with ggg179 marked in red, gg185 marked in purple, and gg193 and gg196 sequences marked in blue.

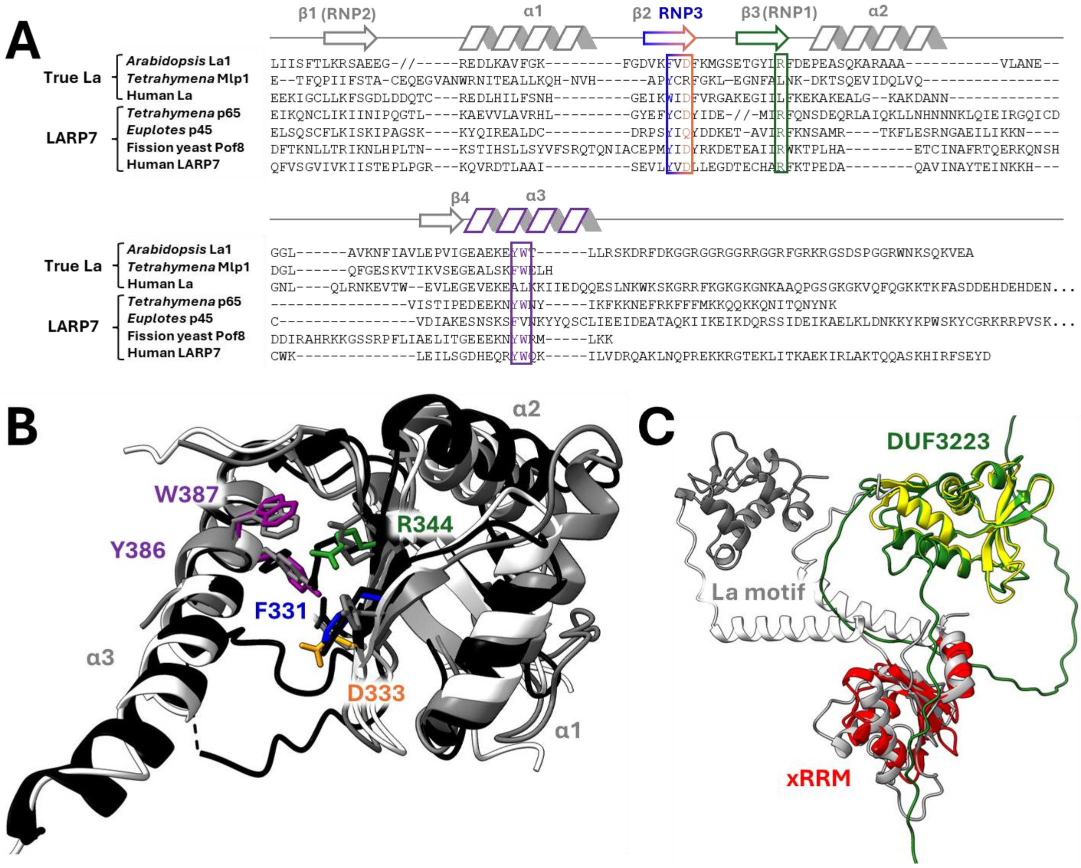

**<u>Figure S5</u> Conservation of xRRM domains. (A)** True La and LARP7 xRRM domain sequence alignment after the style of (7). Annotations show positions of beta sheets (arrows) and alpha helices (zig-zags) with conserved RNA-interacting residues coloured as in Figure 2. The xRRM-specific RNP3 motif is also annotated, as are approximate positions of missing canonical RRM motifs RNP1 and RNP2. **(B)** Alphafold3 (8) prediction of AtLa1 xRRM domain (white) overlaid with experimental models of xRRM domains from fission yeast Pof8 (PDB - 6TZN, grey, 53 paired residues after pruning, RMSD = 0.805 Å) and *Tetrahymena* p65 (PDB - 8GAP, black, 33 paired residues after pruning, RMSD = 0.939 Å ) using the ChimeraX Matchmaker tool (9). Conserved RNA-binding residues are annotated with AtLa1 numbering and colours as in Figure 2. Structural visualisations were prepared using ChimeraX (9). **(C)** Alphafold3 model of *Tetrahymena* Mlp1 showing conserved La motif (grey), predicted xRRM (red) overlaid with predicted AtLa1 xRRM (dark grey), and DUF3223 (yellow) overlaid with predicted AtDomino (green) structures.

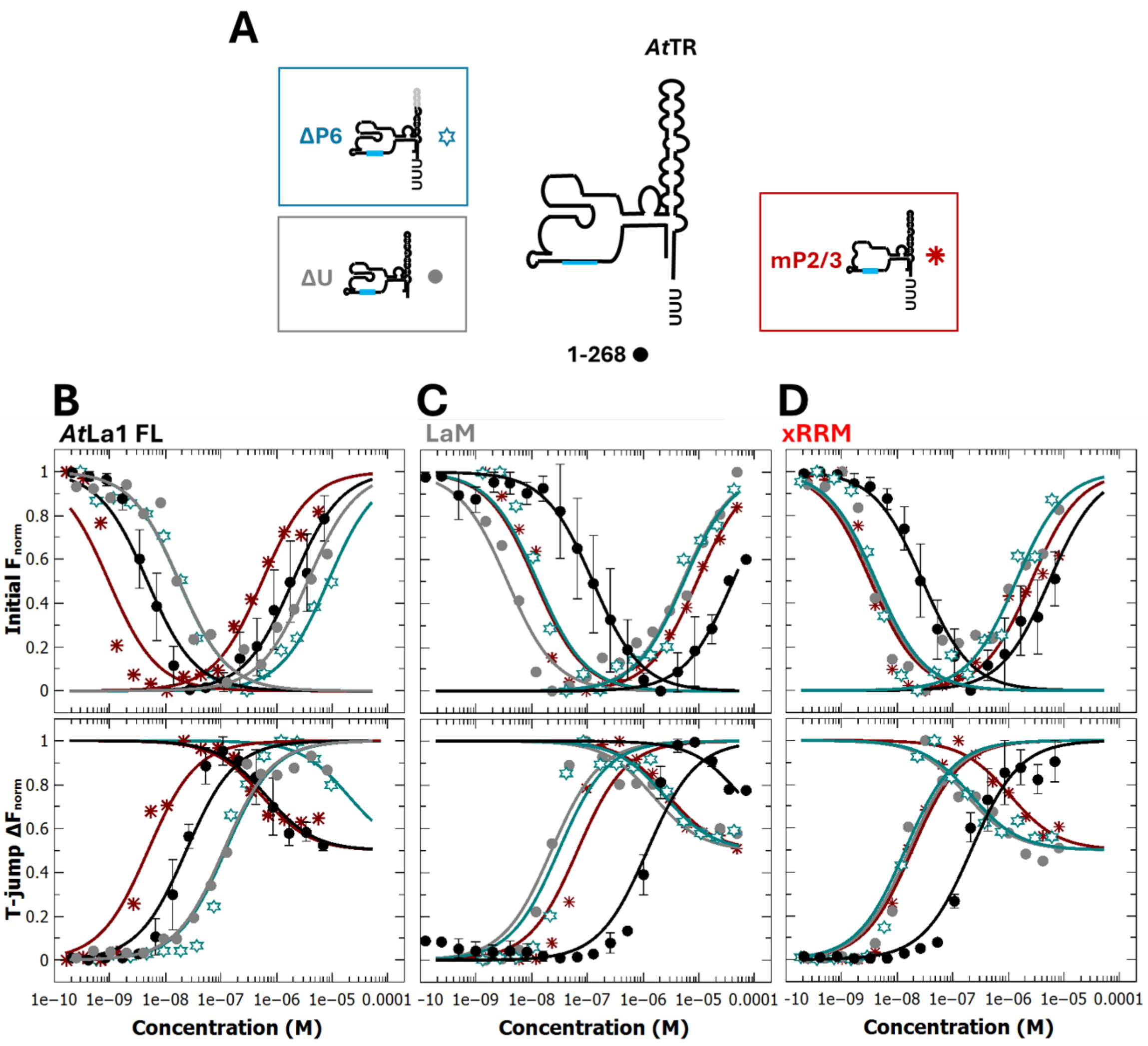

**<u>Figure S6</u> – Extended data of AtLa1-AtTR binding experiments. (A)** Overview of additional AtTR constructs used to map binding of AtLa1 and legend for (B-D). Template region (cyan) and deleted P6 (grey) are highlighted. **(B-D)** Initial capillary fluorescence values from MST experiments (upper panel) or MST 1.5 s T-jump data (lower panel) for 5 nM 3' Cy5-labelled AtTR fragments and La1 FL (B), LaM (C), or xRRM (D) with AtTR 1-268 data reproduced from Figure 2C for comparison.

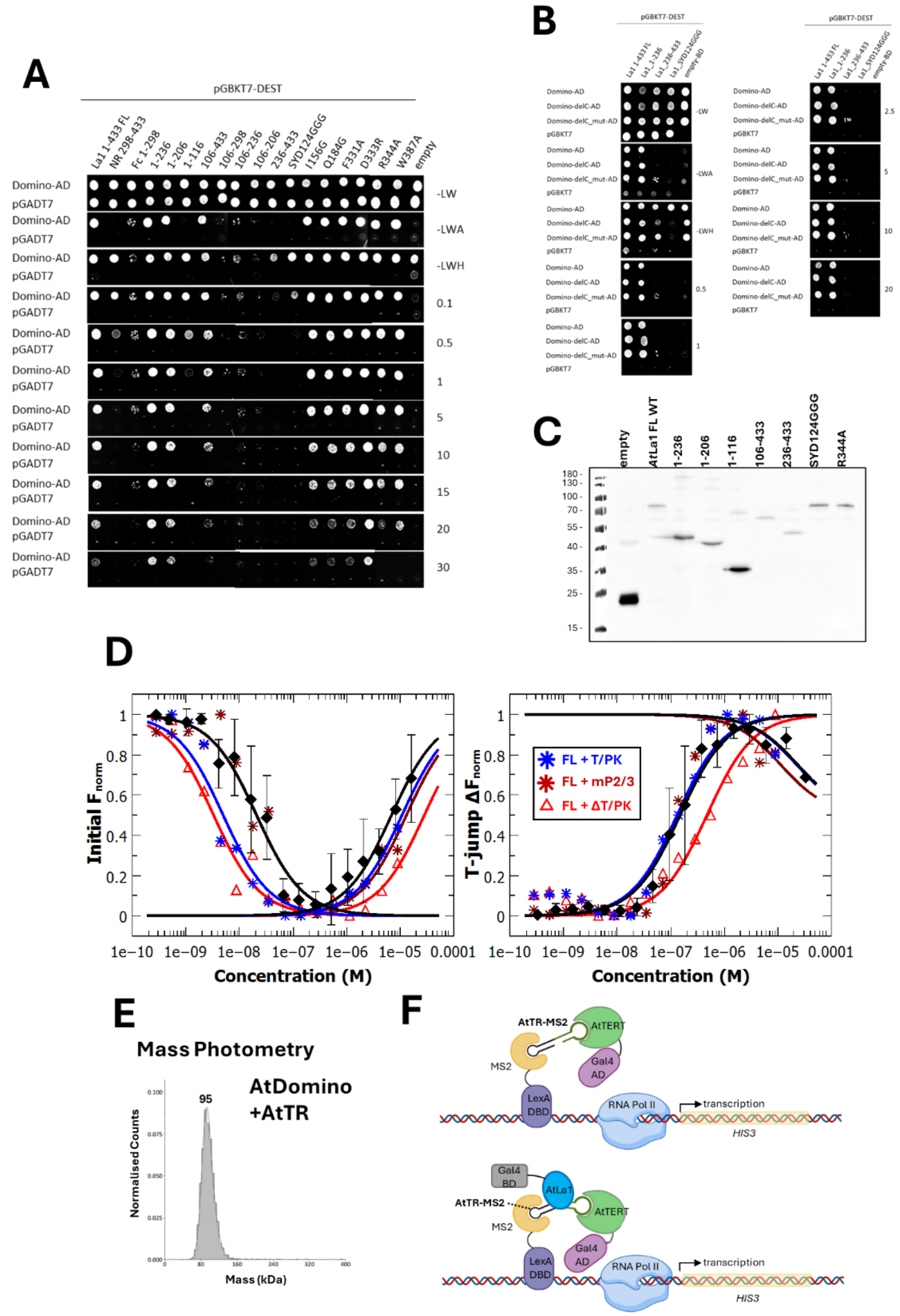

**Figure S7** – **Extended data on protein-protein, RNA-protein and multicomponent protein-RNA interactions. (A-C)** Yeast 2-hybrid assays pairing Gal4 binding domain (BD) fused AtLa1 fragments and variants with Gal4 activation domain (AD) fused (A) AtDomino or (B) AtDomino-delC. Representative anti-BD western blotting of yeast cultures as indicated used for 2-hybrid experiments (C), proteins extracted according to (1). Data shown in Figure 4B is taken from (A). **(D)** Initial capillary fluorescence values (left) and MST 1.5 s T-jump data (right) of AtDomino with 5 nM 3' Cy5-labelled AtTR fragments as annotated. Lines show equations fit to data used to calculate $K_D$, error bars are standard deviation, AtDomino-AtTR 1-268 data are reproduced from Figure 4F. **(E)** Mass photometry deconvoluted mass spectrum of 70 nM AtDomino (predicted 23 kDa) and AtTR 1-262 (82 kDa). Solid lines show Gaussian fittings with values in Table S5, annotated with average mass values in kDa. Single peak is assigned as unresolved AtTR and 1:1 AtTR:AtDomino, with the AtDomino monomer undetectable, due to its low mass **(F)** Schematic of the multicomponent yeast 3-hybrid experiment. Coloured shapes represent proteins, line structures represent RNA (black lines) or DNA (red and blue lines), upper panel is with empty BD and lower panel is in the presence of TR-interacting AtLa1 (blue shape).

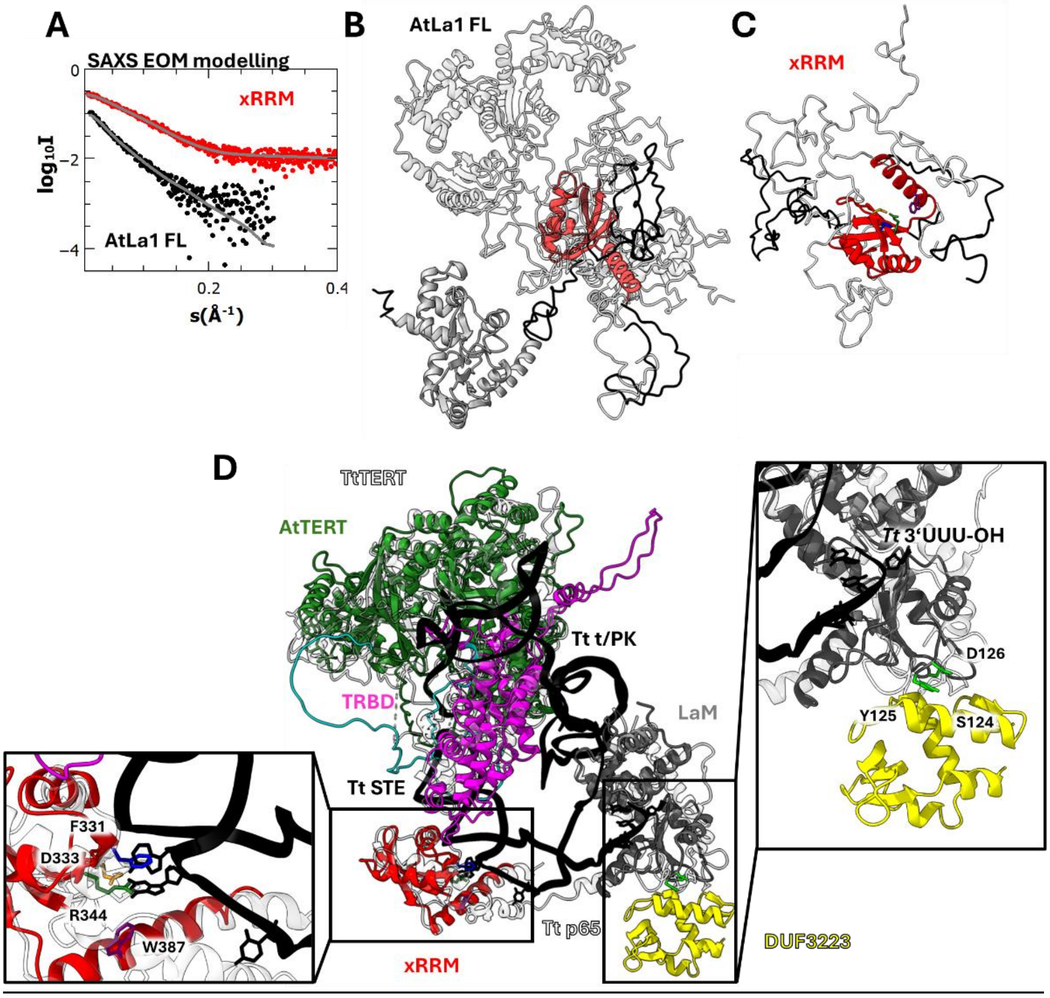

**<u>Figure S8</u> – Overlay of At structural predictions with Tetrahymena telomerase. (A)** Radially averaged small angle X-ray scattering data (dots), fitted to theoretical scattering from structural ensembles (grey lines) calculated by EOM software (2) either AtLa1 FL (black dots, $\chi^2$ = 1.431, SASDYP2) or xRRM (red dots, $\chi^2$ = 1.190, SASDYQ2). **(B,C)** Structural ensembles of (B) AtLa1 FL or (C) xRRM models corresponding to (A). For one model of each ensemble, domains shown are xRRM (red, fixed) and LaM (grey, mobile), plus unstructured linker (black, mobile), with all other models in white. For (C) RNA-implicated residues are shown as sticks and coloured as per Figure 2A. Further details in Table S6 and SASBDB. **(D)** CryoEM derived structural model of Tetrahymena telomerase ((10), 8GAP) with p50 and Teb proteins omitted for clarity, proteins shown as transparent cartoons, telomerase RNA backbone in black with flipped-out residues shown as sticks. Alphafold2 (11) and Alphafold3 (8) predictions of AtLa1 LaM (grey, 72 paired residues after pruning, RSMD = 1.182 Å), xRRM (red, 30 paired residues after pruning, RSMD = 1.038 Å), AtDomino (yellow, predicted complex with AtLa1 for comparison, no equivalent in 8GAP) and AtTERT (green, 273 paired residues after pruning, 1.322 Å) have been overlaid with 8GAP using the ChimeraX tool MatchMaker**,** visualised in ChimeraX (9). Side panels show details of conserved xRRM residues (left) and the semi-conserved loop responsible for interactions between AtLa1 RRM and Domino.

**Table S1. Oligonucleotides used for cloning, mutagenesis, and in PCR for in vitro synthesis**

| Name | note | | sequence (5’-3’) |
|---|---|---|---|
| **Plant TRs** | | | |
| Slycopersicum-R8_Fw | cloning | Eva-C251 | AGGGGGTGTGGAGCTGAGATCAAT |
| Slycopersicum-R8_Rev | cloning, PCR for in vitro RNA synthesis | Eva-C252 | GGAGGTGCTGGCAAGCGAG |
| Cestrum-R8_Fw | cloning | Eva-C253 | CTCGGGGGTGTGGTGTGAGTACTG |
| Cestrum-R8_Rev | cloning, PCR for in vitro RNA synthesis | Eva-C254 | GGGGTGTTTGGCAAGCAAGGAAAC |
| PisumR8_Fw1 | cloning | Eva-C243 | CAGTTTATAGGAGGCTCGCACAACT |
| PisumR8_Rev1 | cloning, PCR for in vitro RNA synthesis | Eva-C244 | CATTATCTAAAATAGTGGTGGGGTATGGG |
| AsparagusR8_Fw1 | cloning | Eva-C245 | AGGGGATGGTGGGCCGTG |
| AsparagusR8_Rev1 | cloning, PCR for in vitro RNA synthesis | Eva-C246 | GGGGGATGGTGTTTTAAGTGAGGC |
| ZmR8_Fw1 | cloning | Eva-C247 | CCAGAGGGTGAAGTGTCATTGGAGT |
| ZmR8_Rev1 | cloning, PCR for in vitro RNA synthesis | Eva-C248 | TGAGGGGGTGCAGCTGGAAG |
| **RNA constructs** | | | |
| U6.26F | cloning | Eva-C180 | AGGGGACATCCGATAAAATTGGAAC |
| U6.26R | cloning, PCR for in vitro RNA synthesis | Eva-C181 | TGCAAAAAAAATTTGGACCATTTCTCGAT |
| MRP1_RNA_Fw | cloning | Eva-C184 | AATTGTCACTGGACGAAGTGAA |
| MRP1_RNA_Rev | cloning, PCR for in vitro RNA synthesis | Eva-C185 | TTGTCATTGAAACGTAAGCCC |
| MRP_RNA_Rev2 | cloning, PCR for in vitro RNA synthesis | Eva-C293 | CGACGAAAGAAAAAGTTGCTTGTCATTGAAACGT |
| ATR-5UTR_Fwclon | cloning | Eva-C177 | AGCGGAAAAACCAGTTAAACCCTT |

| | | | |
|---|---|---|---|
| ATR-5UTR_Rev | cloning, PCR for in vitro RNA synthesis | Eva-C178 | TTCACAATCGCAGAAGCTCTC |
| T7-U6.26F | PCR for in vitro RNA synthesis | Eva-C182 | TAATACGACTCACTATAGGGAGGGGACATCCGATAAAATTGGAAC |
| T7_MRP1_RNA_Fw | PCR for in vitro RNA synthesis | Eva-C186 | TAATACGACTCACTATAGGGCAATTGTCACTGGACGAAGTGAA |
| pCDT1a_T7-1_Fw | PCR for in vitro RNA synthesis | Eva-C135 | TAATACGACTCACTATAGGGTCGTTCAGAGACAGCAATTCATAAT |
| promCDT1a-ATGRev | PCR for in vitro RNA synthesis | Eva-C119 | GTGTACTCATTTTTATCAACAATGTCT |
| F_up_pretsnoR43.1 | cloning | Eva-C300 | CATAGCGTAAGGTTCAACAAAGC |
| R_dw_pretsnoR43.1 | cloning | Eva-C299 | GCAGGCTATTTTGGCAGATAC |
| T7_pretsnoR43.1 | PCR for in vitro RNA synthesis | Eva-C301 | TAATACGACTCACTATAGGGAGCACCAGTGGTCTAGTG |
| R_pU_pretsnoR43.1 | PCR for in vitro RNA synthesis | Eva-C304 | AAAAATAGAGCTCAGAGTAGGC |
| R_pretsnoR43.1 | PCR for in vitro RNA synthesis | Eva-C303 | GAGCTCAGAGTAGGCGAAATC |
| T7_snoR43.1 | PCR for in vitro RNA synthesis | Eva-C302 | TAATACGACTCACTATAGGGTGAGCTGTGATGAAATTGGCTT |
| **Arabidopsis TR mutagenesis** | | | |
| uATR8d201-31 | cloning, delta P6 | Eva-C290 | CGTAGGTGGTTGAAAAACCGTTTCCTCGCCTTACC |
| dATR8d201-31 | cloning, delta P6 | Eva-C289 | GAAACGGTTTTTCAACCACCTACGAACCAAACC |
| U_ATR8_gg179cc | site mutagenesis | Eva-M15 | aagtagaccaggaggttccgtttggttcgtaggtgg |
| d_ATR8_gg179cc | site mutagenesis | Eva-M16 | ccacctacgaaccaaacggaacctcctggtctactt |
| U_ATR8_ggg179cac | site mutagenesis | Eva-M25 | caaaaagtagaccaggaggttcactttggttcgtaggtggttctg |
| d_ATR8_ggg179cac | site mutagenesis | Eva-M26 | cagaaccacctacgaaccaaagtgaacctcctggtctactttttg |
| U_ATR8_gg193cc | site mutagenesis | Eva-M19 | caggaggttgggtttggttcgtacctggttctgttga |
| d_ATR8_gg193cc | site mutagene | Eva-M20 | tcaacagaaccaggtacgaaccaaacccaacctcctg |

| | | | |
|---|---|---|---|
| | sis | | |
| U_ATR8 _gg196cc | site mutagene sis | Eva-M21 | gttgggtttggttcgtaggtccttctgttgaaactagattag |
| d_ATR8 _gg196cc | site mutagene sis | Eva-M22 | ctaatctagtttcaacagaaggacctacgaaccaaacccaac |
| U_ATR8 _gg185cc | site mutagene sis | Eva-M17 | gaccaggaggttgggtttccttcgtaggtggttctg |
| d_ATR8 _gg185cc | site mutagene sis | Eva-M18 | cagaaccacctacgaaggaaacccaacctcctggtc |
| **Arabidopsis TR constructs (PCR for in vitro RNA synthesis)** | | | |
| ATR8_T7-1_Fw | | Eva-C98 | TAATACGACTCACTATAGGGAAGGGGTGTGGGAACCTAG |
| ATR8_T7-26_Fw | | Eva-C99 | TAATACGACTCACTATAGGGAGTCTGCTTATTGATTGC |
| ATR8_T7-178_Fw | | Eva-C100 | TAATACGACTCACTATAGGGTTGGGTTTGGTTCGTAGGTG |
| ATR8_T7-D19-171_Fw | | Eva-C183 | TAATACGACTCACTATAGGGAAGGGGTGTGGGAACCTAAGGAGGTTGGGTTTGGTTCG |
| ATR8_REV1 | | Eva-C86 | AAATATTTGGGGGTGGGAGGGTAA |
| ATR8_152_Rev | | Eva-C103 | AGCCTCTTATGTAGCCATCGA |
| ATR8_251_Rev | | Eva-C102 | AGGGTAAGGCGAGGAAACGGT |
| ATR8_262_Rev | | Eva-C239 | TTTGGGGGTGGGAGGGTAAG |
| ATR8_Rev_Jap | | Eva-C240 | GGGGGTGGGAGGGTAAGG |
| ATR8_245_Rev | | Eva-C288 | AAGGCGAGGAAACGGTTAACC |
| ATR8_dP4_Rev | | dP4_Rev | TTTGGGGGTGGGAGAGGA |
| ATR8-Rev_ter3 | | Eva-C121 | GATGTAAAAACCATCATTACTTAAAATAA |
| ATR8-Rev_ter3B | | Eva-C153 | AAAAACCATCATTACTTAAAATAAAAAATAG |
| ATR8-Rev_ter2 | | Eva-C122 | CTTAAAATAAAAAATAGAAAACAAAATATTTGGG |
| ATR8-Rev_ter2B | | Eva-C154 | AAAATAAAAAATAGAAAACAAAATATTTGGGG |
| ATR8-Rev_ter1 | | Eva-C123 | ATAGAAAACAAAATATTTGGGGGTG |
| ATR8-Rev_ter1B | | Eva-C155 | AAAACAAAATATTTGGGGGTGGGA |
| **universal PCR for in vitro RNA synthesis** | | | |
| M13F | universal | | GTTTTCCCAGTCACGA |

| | | | |
|---|---|---|---|
| **Preparation of Y3H constructs** | | | |
| prBS062 | ΔT/PK and ΔP6 | | GGCTAGAACTAGTGGATCCCCCGGGAAGGGGTGTGGGAACCTAGG |
| prBS063 | ΔT/PK and ΔP6 | | CCTGCAGGCATGCAAGCTGCCCGGGAAATATTTGGGGGTGGGAGGG |
| prBS047 | AtTR1-268 | | CCGGCTAGAACTAGTGGATCCCCCGGGGGGTTTGGTTCGTAGGTGG |
| prBS031 | AtTR1-268 | | CCTGCAGGCATGCAAGCTGCCCGGGAGGGTAAGGCGAGGAAACG |
| **Protein constructs and fragments** | | | |
| F_LA1_GW | | Lada-125 | AAAAAGCAGGCTACATGTCGATTCCTTGTCTAACCG |
| R_LA1+GW | | Lada-126 | AGAAAGCTGGGTCTCATGCTTCCACCTTCTGAGA |
| F_LA1_pGEX4T3 | | Eva-C138 | GTGGATCCCCGAATTCCATGTCGATTCCTTGTCTAACCG |
| R_LA1_pGEX-3 | | Eva-C132 | GTCGACCCGGGAATTTGCTTCCACCTTCTGAGATTTG |
| F-La1_delN_pB1 | | Eva-G218 | AAAAAGCAGGCTACAAAGAAGAAAAGGGTGCTCTAGC |
| R-La1_delC(+)_pB2 | | Eva-G219 | AGAAAGCTGGGTCTTATTTCTCATCATCTTCATCCTCTGC |
| F-La1_G236_pB1 | | Eva-G241 | AAAAAGCAGGCTACGGTTTGATCATTTCATTCACCCTC |
| R-La1_G236(+)_pB2 | | Eva-G242 | AGAAAGCTGGGTCTTAACCCTTGGGATAATCTGGCTCAT |
| R-La1_A206(+)_pB2 | | Eva-G243 | AGAAAGCTGGGTCTTAAGCAAATTTTACTTCATCCTTCTCC |
| R-La1_R116(+)_pB2 | | Eva-G244 | AGAAAGCTGGGTCTTATCTAGCATTAAGTTGCTCAATCAA |
| F-La1_L106_pB1 | | Eva-G245 | AAAAAGCAGGCTACCTGGAAGATTTGATTGAGCAACTT |
| F-DOMINO1_gw | | Eva-G95 | AAAAAGCAGGCTACATGGCTGAAGAACAAGAGATCG |
| R-DOMINO1+gw | | Eva-G96 | AGAAAGCTGGGTCTCATCTTCTGAATCTCCCTCCTC |
| DOMINO1_delC_pB2 | | Eva-G280 | AGAAAGCTGGGTCTTAAGGCAATGGGAGGATTTGATC |
| **La1 mutagenesis** | | | |
| U_La1-F331A | | Eva-M1 | cagatcccatcttgaaatcaacagccttgacatcaccaaattttccga |
| d_La1-F331A | | Eva-M2 | tcggaaaatttggtgatgtcaaggctgttgatttcaagatgggatctg |
| U_La1-D333R | | Eva-M5 | cgtctcagatcccatcttgaaacgaacaaacttgacatcaccaaat |
| d_La1-D333R | | Eva-M6 | atttggtgatgtcaagtttgttcgtttcaagatgggatctgagacg |
| U_La1-R344A | | Eva-M7 | atgcttcaggttcatcaaacgcaagataacccgtctcagatc |
| d_La1-R344A | | Eva-M8 | gatctgagacgggttatcttgcgtttgatgaacctgaagcat |
| U_La1-W387A | | Eva-M9 | tttgcttcgtaggagagtcgcatactccttttcagcttcgc |
| d_La1-W387A | | Eva-M10 | gcgaagctgaaaaggagtatgcgactctcctacgaagcaaa |

| | | | |
|---|---|---|---|
| dLa124SYD-GGG | | Eva-M47 | gattccacgtcttcccttttaacaccaccgccgaaaggtgacgcagctactgttctagc |
| uLa124SYD-GGG | | Eva-M48 | gctagaacagtagctgcgtcacctttcggcggtggtgttaaaagggaagacgtggaatc |
| dLaI156G | | Eva-M49 | tcatgttgcagagtcgaggggattttctggtgttgccttg |
| uLaI156G | | Eva-M50 | caaggcaacaccagaaaatcccctcgactctgcaacatga |
| dLaQ184G | | Eva-M51 | agaatttggtctttgctggtggggagttggagctgaaacc |
| uLaQ184G | | Eva-M52 | ggtttcagctccaactccccaccagcaaagaccaaattct |

**Table S2. $K_D$ values (nM) from ELISA competition experiments.** All experiments used immunodetection of GST-fused La1 FL bound to immobilised 3’ biotinylated AtTR 1-268 and displaced by the presence of unlabelled competitor as indicated.

**Variable Competitor**

| RNA | ELISA | Ratio vs 1-268 | replicas |
|---|---|---|---|
| **AtTR** | 41 ± 6 | 1.00 | 2 |
| **pre-tsnoR43.1** | 150 ± 40 | 3.66 | 2 |
| **pre-tsnoR43.1ΔU** | 100 ± 30 | 2.44 | 2 |
| **snoR43.1** | 14 ± 3 | 0.34 | 2 |
| **snoR43.1 ΔU** | 70 ± 10 | 1.71 | 2 |
| **ytRNA** | 220 ± 30 | 5.37 | 2 |
| **MRP1** | 160 ± 40 | 3.90 | 2 |
| **MRP1 ΔU** | 130 ± 40 | 3.17 | 2 |
| **U6-26** | 260 ± 80 | 6.34 | 2 |
| **ATR 5'UTR** | 600 ± 200 | 14.63 | 2 |

**Variable Competitor**

| AtTR | ELISA | Ratio vs 1-268 | replicas |
|---|---|---|---|
| **1-245** | 9 ± 3 | 0.22 | 2 |
| **1-251** | 50 ± 20 | 1.22 | 2 |
| **1-259** | 52 ± 9 | 1.27 | 2 |
| **0-274** | 90 ± 20 | 2.20 | 2 |
| **0-277** | 70 ± 10 | 1.71 | 2 |
| **0-286** | 90 ± 20 | 2.20 | 2 |
| **0-289** | 25 ± 5 | 0.61 | 2 |
| **0-304** | 13 ± 3 | 0.32 | 2 |
| **0-309** | 32 ± 9 | 0.78 | 2 |
| **T/PK** | 190 ± 40 | 4.63 | 2 |
| **ΔT/PK** | 70 ± 10 | 1.71 | 2 |
| **ΔP4** | 70 ± 10 | 1.71 | 2 |
| **ΔP6** | 80 ± 20 | 1.95 | 2 |
| **P4/5/6** | 500 ± 100 | 12.20 | 2 |
| **ΔU** | 58 ± 9 | 1.41 | 2 |

| **Plant TR Competitor** | Telomere repeat | ELISA | Ratio vs 1-268 | replicas |
|---|---|---|---|---|
| ***Arabidopsis thaliana*** | TTTAGGG | 41 ± 6 | 1.00 | 2 |
| ***Solanum lycopersicum*** | TTTAGGG | 30 ± 10 | 0.73 | 2 |
| ***Asparagus officinalis*** | TTAGGG | 50 ± 10 | 1.22 | 2 |
| ***Allium cepa*** | CTCGGTTATGGG | 50 ± 10 | 1.22 | 2 |
| ***Nicotiana sylvestris*** | TTTAGGG | 42 ± 9 | 1.02 | 2 |
| ***Scilla peruviana*** | TTAGGG | 80 ± 30 | 1.95 | 2 |
| ***Zea mays*** | TTTAGGG | 60 ± 20 | 1.46 | 2 |
| ***Cestrum elegans*** | TTTTTTAGGG | 90 ± 20 | 2.20 | 2 |
| ***Pisum sativum*** | TTTAGGG | 80 ± 20 | 1.95 | 2 |

**Table S3. Full details of $K_D$ values from MST experiments.** Processes Fluor 1 and Fluor 2 are fast and slow initial fluorescence changes. MST values are calculated from the first 1.5 s of thermophoresis (T-jump) fast (1) and slow (2) processes. Error values are from $K_D$ fittings, some data are replicated from Table 1 for completeness.

| | | ***K<sub>D</sub>* at 25 °C** | | | | **Ratio vs AtTR 1-268** | | | | |
|---|---|---|---|---|---|---|---|---|---|---|
| Protein | AtTR | Fluor 1 (nM) | Fluor 2 (µM) | MST 1 (nM) | MST 2(µM) | Fluor 1 | Fluor 2 | MST 1 | MST2 | replicas |
| **AtLa1 FL** | **1-268** | 5 ± 1 | 1.9 ± 0.2 | 20 ± 10 | 0.65 ± 0.1 | 1.0 | 1.0 | 1.0 | 1.0 | 5 |
| | **mP2/3** | 1 ± 0.3 | 0.54 ± 0.1 | 5 ± 2 | 0.5 ± 0.1 | 0.2 | 0.3 | 0.3 | 0.8 | 1 |
| | **T/PK** | 21 ± 2 | 90 ± 30 | 1000 ± 200 | | 4.2 | 47.4 | 50.0 | | 1 |
| | **ΔT/PK** | 2.7 ± 0.7 | 1.4 ± 0.1 | 16 ± 7 | 1.4 ± 0.2 | 0.5 | 0.7 | 0.8 | 2.2 | 1 |
| | **ΔP4** | 38 ± 4 | 35 ± 6 | 560 ± 70 | | 7.6 | 18.4 | 28.0 | | 1 |
| | **ΔP6** | 16 ± 6 | 8.1 ± 0.7 | 130 ± 30 | | 3.2 | 4.3 | 6.5 | | 1 |
| | **P4/5/6** | 6 ± 3 | 3 ± 0.2 | 34 ± 9 | 4 ± 0.2 | 1.2 | 1.6 | 1.7 | 6.2 | 1 |
| | **ΔU** | 12 ± 2 | 2.4 ± 0.3 | 70 ± 10 | | 2.4 | 1.3 | 3.5 | | 2 |
| **AtLa1 LaM** | **1-268** | 130 ± 10 | 41 ± 3 | 1200 ± 300 | 70 ± 30 | 1.0 | 1.0 | 1.0 | 1.0 | 3 |
| | **mP2/3** | 12 ± 3 | 9.4 ± 0.3 | 70 ± 40 | 2.6 ± 0.3 | 0.1 | 0.2 | 0.1 | 4.0 | 1 |
| | **T/PK** | 5 ± 1 | 10 ± 1 | 170 ± 20 | | 0.0 | 0.2 | 0.1 | | 1 |
| | **ΔT/PK** | 340 ± 70 | 140 ± 30 | 2000 ± 500 | | 2.6 | 3.4 | 1.7 | | 1 |
| | **ΔP4** | 170 ± 40 | 80 ± 10 | 2100 ± 300 | | 1.3 | 2.0 | 1.8 | | 1 |
| | **ΔP6** | 13 ± 3 | 5.6 ± 0.9 | 30 ± 20 | 3.1 ± 0.7 | 0.1 | 0.1 | 0.0 | 4.8 | 1 |
| | **P4/5/6** | 130 ± 30 | 8 ± 2 | 600 ± 100 | 18 ± 4 | 1.0 | 0.2 | 0.5 | 27.7 | 1 |
| | **ΔU** | 4 ± 1 | 5.6 ± 0.8 | 20 ± 10 | 2.6 ± 0.4 | 0.0 | 0.1 | 0.0 | 4.0 | 1 |
| **AtLa1 xRRM** | **1-268** | 28 ± 3 | 6 ± 1 | 220 ± 30 | | 1.0 | 1.0 | 1.0 | | 2 |
| | **mP2/3** | 3 ± 1 | 2.6 ± 0.5 | 19 ± 8 | 1 ± 0.1 | 0.1 | 0.4 | 0.1 | 1.5 | 1 |
| | **T/PK** | 18 ± 5 | 30 ± 5 | 590 ± 90 | | 0.6 | 5.0 | 2.7 | | 1 |
| | **ΔT/PK** | 4 ± 1 | 2.3 ± 0.2 | 28 ± 5 | 1.4 ± 0.6 | 0.1 | 0.4 | 0.1 | 2.2 | 1 |
| | **ΔP4** | 4 ± 1 | 2.4 ± 0.3 | 40 ± 10 | 2.1 ± 0.8 | 0.1 | 0.4 | 0.2 | 3.2 | 1 |
| | **ΔP6** | 4 ± 1 | 1.4 ± 0.2 | 14 ± 7 | 1 ± 0.7 | 0.1 | 0.2 | 0.1 | 1.5 | 1 |
| | **P4/5/6** | 18 ± 5 | 7.6 ± 0.9 | 90 ± 30 | 8 ± 1 | 0.6 | 1.3 | 0.4 | 12.3 | 1 |
| | **ΔU** | 4 ± 1 | 1.4 ± 0.3 | 20 ± 10 | 0.18 ± 0 | 0.1 | 0.2 | 0.1 | 0.3 | 1 |
| **AtLa1 F331A** | **1-268** | 8 ± 2 | 1.8 ± 0.3 | 24 ± 3 | 3 ± 1 | | | | | 4 |
| **AtLa1 D333R** | **1-268** | 3 ± 1 | 1.2 ± 0.1 | 29 ± 4 | 0.45 ± 0.1 | | | | | 3 |
| **AtLa1 R344A** | **1-268** | 48 ± 7 | 14 ± 4 | 550 ± 60 | | | | | | 2 |
| **AtLa1 W387A** | **1-268** | 40 ± 4 | 2.7 ± 0.9 | 230 ± 50 | 4 ± 1 | | | | | 2 |
| **AtLa1 FL** | **gg179cc** | 2.2 ± 0.4 | 0.9 ± 0.3 | 15 ± 4 | 1.5 ± 0.2 | 0.4 | 0.5 | 0.8 | 2.3 | 1 |
| | **ggg179cac** | 3 ± 0.7 | 3.2 ± 0.3 | 31 ± 5 | 1.8 ± 0.5 | 0.6 | 1.7 | 1.6 | 2.8 | 1 |
| | **gg185cc** | 18 ± 4 | 4.6 ± 0.6 | 140 ± 30 | 1.2 ± 0.2 | 3.6 | 2.4 | 7.0 | 1.8 | 1 |
| | **gg193cc** | 3.2 ± 0.6 | 1.3 ± 0.3 | 19 ± 3 | 0.6 ± 0.2 | 0.6 | 0.7 | 1.0 | 0.9 | 1 |
| | **gg196cc** | 3 ± 0.7 | 2.9 ± 0.3 | 49 ± 8 | 4 ± 1 | 0.6 | 1.5 | 2.5 | 6.2 | 1 |
| **AtLa1 FL** | **pre-tsnoR43.1** | 20 ± 3 | 6.8 ± 0.7 | 240 ± 30 | | 4.0 | 3.6 | 12.0 | | 2 |
| | **pre-tsnoR43.1 ΔU** | 10 ± 3 | 10 ± 3 | 180 ± 20 | | 2.0 | 5.3 | 9.0 | | 2 |
| | **tsnoR43.1** | 1.3 ± 0.4 | 5.6 ± 0.6 | 50 ± 5 | 4 ± 2 | 0.3 | 2.9 | 2.5 | 6.2 | 1 |
| | **tsnoR43.1 ΔU** | 40 ± 10 | 29 ± 7 | 1000 ± 90 | | 8.0 | 15.3 | 50.0 | | 1 |
| | **ytRNA** | 100 ± 30 | 17 ± 8 | 1300 ± 200 | | 20.0 | 8.9 | 65.0 | | 2 |
| | **MRP1** | 3 ± 0.7 | 2.7 ± 0.5 | 120 ± 30 | 8 ± 4 | 0.6 | 1.4 | 6.0 | 12.3 | 1 |

| | | | | | | | | | | |
|---|---|---|---|---|---|---|---|---|---|---|
| | **MRP1 ΔU** | 5 ± 2 | 1.6 ± 0.4 | 53 ± 7 | 9 ± 3 | 1.0 | 0.8 | 2.7 | 13.8 | 1 |
| **TRBD-linker** | **1-268** | 8 ± 2 | 3.9 ± 0.3 | 39 ± 5 | 3.4 ± 0.6 | | | | | 3 |
| **TRBD** | **1-268** | 8 ± 2 | 7 ± 1 | 42 ± 6 | 8 ± 0.9 | | | | | 2 |
| **TRBD-trunc** | **1-268** | 13 ± 2 | 110 ± 20 | 710 ± 70 | | | | | | 1 |
| **Domino FL** | **1-268** | 21 ± 3 | 6.9 ± 0.7 | 150 ± 20 | 24 ± 6 | 1.0 | 1.0 | 1.0 | 1.0 | 3 |
| | **mP2/3** | 21 ± 7 | 14 ± 2 | 150 ± 30 | | 1.0 | 2.0 | 1.0 | | 1 |
| | **T/PK** | 5 ± 1 | 10.9 ± 0.4 | 130 ± 20 | | 0.2 | 1.6 | 0.9 | | 1 |
| | **ΔT/PK** | 3.3 ± 0.6 | 29 ± 5 | 470 ± 60 | | 0.2 | 4.2 | 3.1 | | 1 |
| | **ytRNA** | 1400 ± 600 | 70 ± 50 | 3700 ± 600 | | 66.7 | 10.1 | 24.7 | | 1 |
| **Domino delC** | **1-268** | 90 ± 20 | 6 ± 2 | 220 ± 50 | 8 ± 4 | | | | | 1 |

**Table S4. $K_D$ values from MST experiments with competitors.** 'Additive' indicates a constant concentration of non-labelled RNA competitor as indicated or the presence of another protein as indicated. Processes Fluor 1 and Fluor 2 are fast and slow initial fluorescence changes. MST values are calculated from the first 1.5 s of thermophoresis (T-jump) fast (1) and slow (2) processes. Error values are from $K_D$ fittings.

| | | | | | | **Ratio vs AtTR 1-268** | | | | |
|---|---|---|---|---|---|---|---|---|---|---|
| Protein +Constant Competitor | Additive Competitor | Fluor 1 (nM) | Fluor 2 (µM) | MST 1 (nM) | MST2 (µM) | Fluor1 | Fluor 2 | MST 1 | MST2 | replicas |
| **FL+TR** | **ytRNA (100x)** | 9 ± 2 | 0.7 ± 0.2 | 310 ± 30 | | 1.8 | 0.4 | 15.5 | | 1 |
| | **ytRNA (1000x)** | 30 ± 8 | 0.9 ± 0.3 | 530 ± 70 | | 6.0 | 0.5 | 26.5 | | 1 |
| | **pre-tsnoR43.1 (10x)** | 60 ± 30 | | 700 ± 100 | | 12.0 | | 35.0 | | 1 |
| | **pre-tsnoR43.1 ΔU (10x)** | 110 ± 30 | | 900 ± 100 | | 22.0 | | 45.0 | | 1 |
| **LaM+TR** | **ytRNA (100x)** | 130 ± 40 | 80 ± 20 | 1100 ± 100 | | 1.0 | 2.0 | 0.9 | | 1 |
| **xRRM +TR** | **ytRNA (100x)** | 18 ± 2 | 6 ± 1 | 250 ± 40 | | 0.6 | 1.0 | 1.1 | | 1 |
| **TRBD+TR** | **La1 (125 nM)** | | 0.25 ± 0.1 | | | | | | | 1 |
| | **La1 (1250 nM)** | | 0.17 ± 0.1 | | | | | | | 1 |
| | **xRRM (125 nM)** | | 0.6 ± 0.1 | 25 ± 3 | 0.6 ± 0.3 | | 0.1 | 0.6 | 0.1 | 1 |
| **Domino+TR** | **ytRNA (1000x)** | 5 ± 2 | 3 ± 1 | 690 ± 70 | | 0.2 | 0.4 | 4.6 | | 1 |
| | **La1 (125 nM)** | | 2.7 ± 0.6 | 200 ± 50 | | | 0.4 | 1.3 | | 1 |
| | **La1 (1250 nM)** | | 0.9 ±0.3 | 130 ± 90 | | | 0.1 | 0.9 | | 1 |
| | **xRRM (63 nM)** | | 0.6 ± 0.1 | 180 ± 40 | | | 0.1 | 1.2 | | 1 |
| | **SYD (63 nM)** | 8 ± 2 | 3.6 ± 0.8 | 220 ± 20 | 40 ± 20 | 0.4 | 0.5 | 1.5 | 1.7 | 1 |

**Table S5. Gaussian fittings to deconvoluted mass photometry spectra.** Peak numberings are from low to high mass, σ indicates peak width, % is the percentage of particles covered by the fitting. * indicates that samples were prepared in high concentration as noted and an unknown tiny amount was transferred to the final drop using the so-called tip method (see Methods).

| Sample | Peak1 (kDa) | σ | % | Peak2 (kDa) | σ | % | Peak3 (kDa) | σ | % | Peak4 (kDa) | σ | % | Peak5 (kDa) | σ | % |
|---|---|---|---|---|---|---|---|---|---|---|---|---|---|---|---|
| **AtLa1 (5 nM)** | 55 | 10.4 | 82 | 82 | 14.6 | 17 | 117 | 9.6 | 4 | | | | | | |
| **AtLa1 (10 µM) +TR (10 µM)*** | 51 | 9.2 | 65 | 74 | 11.7 | 25 | 101 | 17.5 | 14 | 138 | 21 | 8 | | | |
| **AtLa1 (5 nM) +Domino (50 nM)** | 53 | 11.3 | 65 | 76 | 13.4 | 41 | 112 | 11.7 | 7 | 137 | 20 | 7 | | | |
| **AtLa1 (10 µM ) +Domino (10 µM ) +TR (10 µM)*** | 56 | 10.2 | 55 | 76 | 16 | 38 | 105 | 19.6 | 21 | 142 | 9.6 | 7 | 177 | 21 | 7 |
| **AtDomino (70 nM) +TR (70 nM)** | 95 | 13.3 | 97 | | | | | | | | | | | | |

**Table S6.** **Summary of integral SAXS structural parameters.**

| Sample code | ***SASDYQ2*** **(xRRM)** | ***SASDYP2*** **(AtLa1)** |
|---|---|---|
| **Data Collection Parameters** | | |
| Instrument | BioSAXS-2000 | BioSAXS-2000 |
| Wavelength [Å] | 1.54 | 1.54 |
| q range [Å$^{1}$] | 0.009 – 0.65 | 0.009 – 0.65 |
| Exposure time [min] | 60 | 60 |
| Temperature [°C] | 20 | 9 |
| pH | 8.0 | 8.0 |
| Concentration [mg ml$^{-1}$] | 1.0 | 3.0 |
| **Structural parameters** | | |
| $R_g$ [Å] (from Guinier) | 26.14 | 49.52 |
| $R_g$ [Å] (from P(r)) | 28.59 | 44.79 |
| $D_{max}$ [Å] | 107.2 | 132.6 |
| Porod volume estimate [Å$^{3}$] | 24157 | 110848 |
| $M_W$ from sequence [kDa] | 22.1 | 48.1 |
| Primus $M_W$ estimation [kDa] | 12 | 94.2 |
